# The chromodomain proteins, Cbx1 and Cbx2 have distinct roles in the regulation of heterochromatin and virulence in the fungal wheat pathogen, *Zymoseptoria tritici*

**DOI:** 10.1101/2022.09.21.508279

**Authors:** Callum J. Fraser, Julian C. Rutherford, Jason J. Rudd, Simon K. Whitehall

## Abstract

Heterochromatin is characterized by specific histone post-translational modifications such as the di- and tri-methylation of histone H3 on lysine 9 (H3K9me2/3), which direct the recruitment of ‘reader’ proteins to chromatin. In the fungal phytopathogen, *Zymoseptoria tritici,* deletion of the H3K9 methyltransferase gene *kmt1,* results in a global increase in the expression of transposable elements (TEs), genome instability and loss of virulence. Here we have identified two *Z. tritici* chromodomain proteins, Cbx1 and Cbx2, that recognise H3K9me modifications. Cbx1 is a Heterochromatin Protein 1 homolog that binds H3K9me2/3 *in vitro* and associates with heterochromatic loci *in vivo*. Transcriptomic analysis also indicates that Cbx1 and Kmt1 regulate overlapping sets of protein-encoding genes. However, unlike Δ*kmt1* mutants, Δ*cbx1* strains do not exhibit a global increase in TE expression and have only a partial reduction in virulence, suggesting the existence of additional H3K9me reader proteins. Accordingly, we have identified a fungal-specific chromodomain protein, Cbx2, that binds H3K9me3 *in vitro*. Strikingly, the growth defects of Δ*cbx1* Δ*cbx2* double mutants closely resemble those of Δ*kmt1* consistent with Cbx1 and Cbx2 playing redundant roles in gene silencing. Overall, the data suggest that key functions of H3K9me modifications are mediated by a combination of Cbx1 and Cbx2.

## INTRODUCTION

*Zymoseptoria tritici* is a fungal pathogen of wheat that is responsible for septoria tritici blotch disease. The initial phase of infection (10-14 days) is characterised by symptomless intercellular colonisation of the stomatal cavity and evasion from detection through the secretion of chitin binding proteins and likely, factors that repress and manipulate the wheat immune response (Goodwin *et al*., 2011, Lee *et al*., 2014, Marshall *et al*., 2011, Rudd, 2015, Steinberg, 2015, Canzio *et al*., 2014, Kumar & Kono, 2020). The second stage of infection is marked by death of the plant cells lining the stomatal cavity and a switch to a necrotrophic growth phase (Rudd, 2015, Steinberg, 2015). The increase in nutrient availability allows a rapid increase in growth followed by the formation of pycnidia, the asexual fruiting bodies of *Z. tritici*, which appear as melanised black dots on the leaf surface (Steinberg, 2015). Unsurprisingly, the switch in lifestyle during the infection process is accompanied by a major reprogramming of the transcriptome (Kellner *et al*., 2014, Rudd *et al*., 2015), but the mechanisms by which this is achieved are poorly understood.

The colonization of plant tissue by fungal pathogens requires the expression of specific effector genes (Uhse & Djamei, 2018). Effector genes are often lowly expressed in axenic culture but are strongly upregulated during the infection process. In a number of plant-associated fungi, putative effector genes are located in heterochromatic regions of the genome that are often enriched with transposable elements (TEs) and subject to transcriptional silencing (Soyer *et al*., 2015). Accordingly, the disruption of heterochromatin in the oil seed rape pathogen *Leptosphaeria maculuns* results in the de-repression of effector genes located in repeat rich regions (Soyer *et al*., 2014) and the expression of genes that allow *Epichloe festucae* to form a mutualistic interaction with the grass species, *Lolium perenne* are also regulated by heterochromatin (Chujo & Scott, 2014). These findings have led to a model whereby reprogramming of heterochromatic regions of the genome regulates effector gene expression programmes and facilitates plant colonization (Soyer *et al*., 2015).

Heterochromatin is characterised by specific histone post translation modifications (PTMs), notably di- and tri-methylation of histone H3 on lysine 9 (H3K9me2/3) and the tri-methylation of lysine 27 (H3K27me/3) (Allshire & Madhani, 2018). H3K9me2/3 is considered to be a hallmark of constitutive heterochromatin whereas H3K27me3 in metazoans is commonly associated with facultative heterochromatin that is reversible in response to appropriate stimuli (Allshire & Madhani, 2018). Genome-wide mapping of these modifications in *Z. tritici* has revealed that H3K9me2 is predominantly associated with TEs (Schotanus *et al*., 2015). H3K27me3 is associated with TEs but is also enriched at telomeres and on the conditionally dispensable accessory chromosomes (Schotanus *et al*., 2015). Deletion of the H3K27 methyltransferase gene, *kmt6* results in increased expression of genes located on accessory chromosomes but has only a subtle impact on virulence in wheat infection assays (Möller *et al*., 2019). In contrast, deletion of *kmt1*, which encodes the H3K9 methyltransferase, results in growth defects *in vitro* and severely compromises virulence (Möller *et al*., 2019). H3K9me2/3 also plays a key role in maintaining genome stability. In Δ*kmt1* strains, loci that were previously occupied by H3K9me3 are invaded by H3K27me3 which is accompanied by an increased frequency of large scale chromosomal rearrangements and accessory chromosome loss (Möller *et al*., 2019).

Histone PTMs modulate chromatin function by directing the recruitment of non-histone proteins ‘reader’ proteins. Recognition of H3K9me2/3 is commonly achieved by members of the Heterochromatin Protein 1 (HP1) family that have a conserved domain organisation comprised of an N-terminal chromodomain (CD), and a C- terminal chromoshadow domain (CSD) separated by a flexible hinge region (Canzio *et al*., 2014, Kumar & Kono, 2020). The CD is responsible for the recognition of H3K9me2/3 whereas the CSD mediates homodimerization and provides a hub for the docking of interacting proteins (Bannister *et al*., 2001, Cowieson *et al*., 2000, Smothers & Henikoff, 2000). An HP1 dimer is capable of bridging two nucleosomes (Machida *et al*., 2018) and it is proposed that CD-CD interactions drive the formation of oligomeric structures which condense chromatin and provide a platform for the assembly of additional heterochromatin components (Canzio *et al*., 2014, Kumar & Kono, 2020).

Here we have identified and characterised the *Z. tritici* HP1 homolog, Cbx1. We find that Cbx1 and the H3K9 methyltransferase, Kmt1 regulate the expression of highly similar sets of protein encoding genes and that Cbx1 is enriched at H3K9me-marked loci. However, the removal of Cbx1 does not result in the phenotypes that are associated with loss of Kmt1, suggesting that *Z. tritici* has additional H3K9me- effectors. Consistent with this hypothesis, we show that a fungal-specific CD protein, Cbx2, binds to H3K9me3 *in vitro* and plays a role in the silencing of some Kmt1- regulated genes. Furthermore, genetic analysis is consistent with a model whereby key biological effects of H3K9me PTMs in *Z. tritici* are mediated by a combination of Cbx1 and Cbx2.

## RESULTS

### Cbx1 is a *Z. tritici* HP1 homolog that binds H3K9me2/3

Methylation of histone H3 on lysine 9 is required for the genome stability and virulence of *Z. tritici* (Möller *et al*., 2019). Therefore, we sought to identify the ‘reader’ proteins that recognise this histone modification. In many organisms, H3K9me2/3 marks are bound by members of the HP1 family of Chromobox (Cbx) proteins. BLAST analyses of the *Z. tritici* genome sequence revealed a hypothetical protein ZtRRes_04004, (hereafter called Cbx1) with an N-terminal chromodomain (CD) and a C-terminal chromoshadow domain (CSD) that share high similarity with HP1 proteins from other fungi (Fig 1). The Cbx1 CD is flanked by an acidic N-terminal patch and basic C- terminal hinge region which are also characteristics of HP1-type proteins (Hiragami-Hamada & Nakayama, 2019).

**Figure 1.**
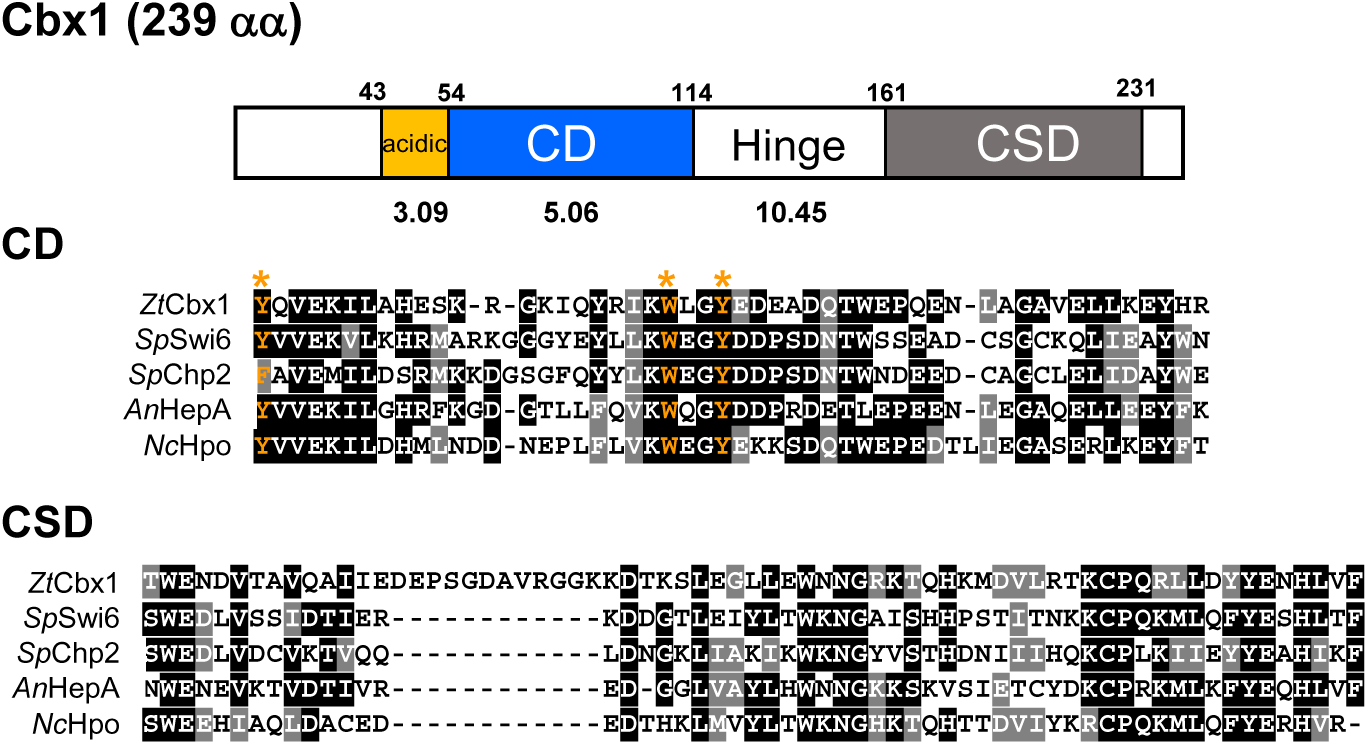
Cbx1 is an HP1 homolog. Schematic representation of the domain architecture of Cbx1 showing the locations and theoretical pI values of the acidic N-terminal patch, chromodomain (CD), hinge, and chromoshadow domain (CSD) (top panel). Sequence alignment of the CD and CSD regions from the indicated fungal HP1 proteins was generated using CLUSTAL (middle and bottom panels). Full shading (black) represents conservation of an amino acid in at least 50% of the sequences, whilst grey shading denotes conservation of a residue of similar chemistry in at least 50% of the analysed sequences. Aromatic ‘methyl-lysine cage’ residues are coloured yellow and their positions are highlighted with asterisks.

While HP1 proteins typically exhibit specificity for H3K9me2/3, both *Tetrahymena* Hhp1 and *Arabidopsis* TFL2/LHP1 recognise H3K27me3 (Turck *et al*., 2007, Yale *et al*., 2016). This prompted us to assess the binding specificity of Cbx1. Full-length Cbx1 was expressed as a GST fusion protein in *E. coli* and purified by affinity and size exclusion chromatography. The binding capacity of GST-Cbx1 was then investigated using a pull-down assay with biotinylated histone H3 peptides. In these assays GST-Cbx1 exhibited a clear preference for H3K9me2 and H3K9me3 modified peptides. No preference for the H3K27me3 or H3K4me3 peptides relative to the unmodified H3 peptide control was observed (Fig 2A & B). These results indicate that Cbx1 is an HP1 family member that binds to H3K9me2/3 modifications *in vitro*.

**Figure 2.**
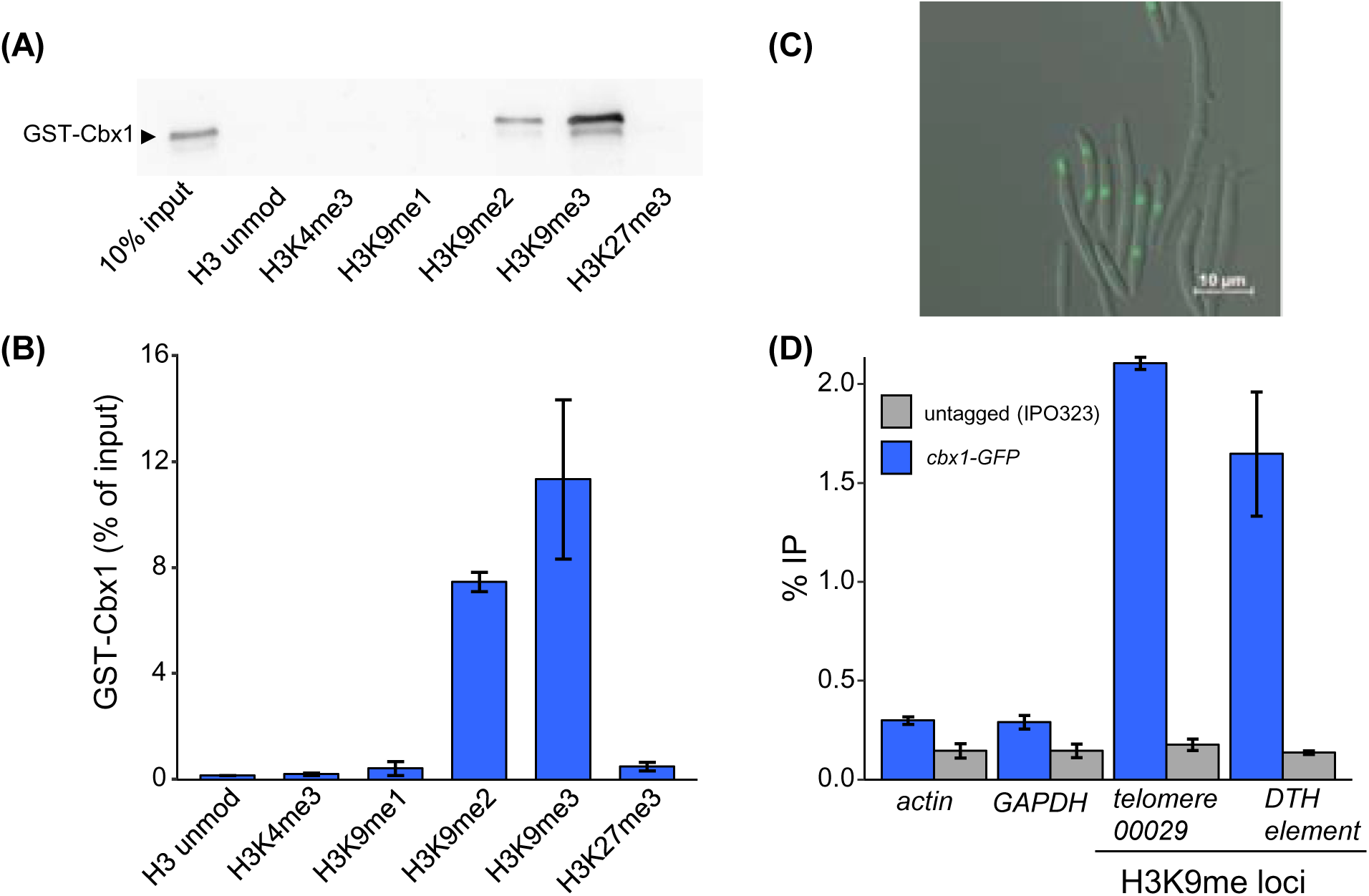
Cbx1 binds H3K9me2/3. **(A)** GST-Cbx1 (1 μg) was incubated with the indicated biotinylated histone H3 peptide and streptavidin beads. Beads were recovered and co-precipitation of GST-Cbx1 was analyzed by western blotting using a GST specific antibody. A 10% input GST-Cbx1 was included as a reference. A representative of three biological repeats is shown. **(B)** Quantification of the GST-Cbx1 signal was relative to the 10% input. Data is the mean of three biological repeats and error bars are ±SEM. **(C)** Fluorescence microscopy of the *cbx1-GFP* strain. **(D)** Cbx1 is associated with H3K9me3 modified chromatin. Chromatin immunoprecipitation (ChIP)-qPCR was used to determine the enrichment of Cbx1-GFP at the indicated loci. The reference IPO323 (untagged) strain was included as a control. %IP was quantified relative to the input sample. Data is the mean of three biological repeats error bars are ±SEM.

The subcellular localisation of Cbx1 was determined by constructing a strain expressing a GFP-tagged fusion protein (*cbx1-GFP*) under the control of its own promoter. Fluorescence microscopy of *cbx1-GFP* cells revealed a strong nuclear GFP signal (Fig 2C). Next, chromatin immunoprecipitation (ChIP) assays were used to investigate the ability of Cbx1 to associate with H3K9me-enriched regions of the genome (Fig 2D). A strong enrichment of Cbx1-GFP was observed at a TE (DTH_element 299 5_ZTIPO323) that is known to be associated with H3K9me (Schotanus *et al*., 2015). Furthermore, a similar enrichment of Cbx1-GFP was also found at a H3K9me-marked subtelomeric region (Chromosome 1: 161011-175642). Importantly, Cbx1-GFP enrichment was not detected at the euchromatic (H3K4me3- associated) genes, *actin* (*Mycgr3G105948*) and *GAPDH* (*Mycgr3G99044*). Notably, *GAPDH*, is located adjacent to an H3K9me-marked DNA transposon, suggesting that the resolution of the assay was sufficient to distinguish between neighbouring H3K9me-marked and non-marked genes. Taken together the data indicate that Cbx1 is an HP1 ortholog and is likely to function in the recognition of H3K9me2/3 in *Z. tritici*.

### Deletion of *cbx1* and *kmt1* results in distinct effects on growth *in vitro* and *in planta*

The role of HP1 in the fitness of *Z. tritici* was investigated by generating *cbx1* deletion strains. For comparison, we also constructed strains lacking the H3K9 methyltransferase gene, *kmt1* and confirmed the loss of H3K9me3 marks in these mutants (Fig S1A). Initially, the *in vitro* growth and stress-sensitivity profiles of the Δ*cbx1* and Δ*kmt1* mutants relative to the IPO323 reference strain were determined. As previously reported, deletion of *kmt1* resulted in a slow growth phenotype (Möller *et al*., 2019) but surprisingly, the Δ*cbx1* strains showed no marked reduction in fitness (Fig 3). Furthermore, although Δ*kmt1* strains were sensitive to osmotic stresses (NaCl, sorbitol), oxidative stress (H_2_O_2_) and cell wall damaging agents (Calcofluor and Congo Red) loss of *cbx1* did not result increase the sensitivity of *Z. tritici* to any of these agents. However, we did note that Δ*cbx1* strains exhibited a slight increase in sensitivity to hydroxyurea which results in reduced dNTP levels and replication stress. Deletion of *cbx1* also resulted in increased levels of melanisation on PD agar (PDA) at 25°C (Fig. 3).

**Figure 3.**
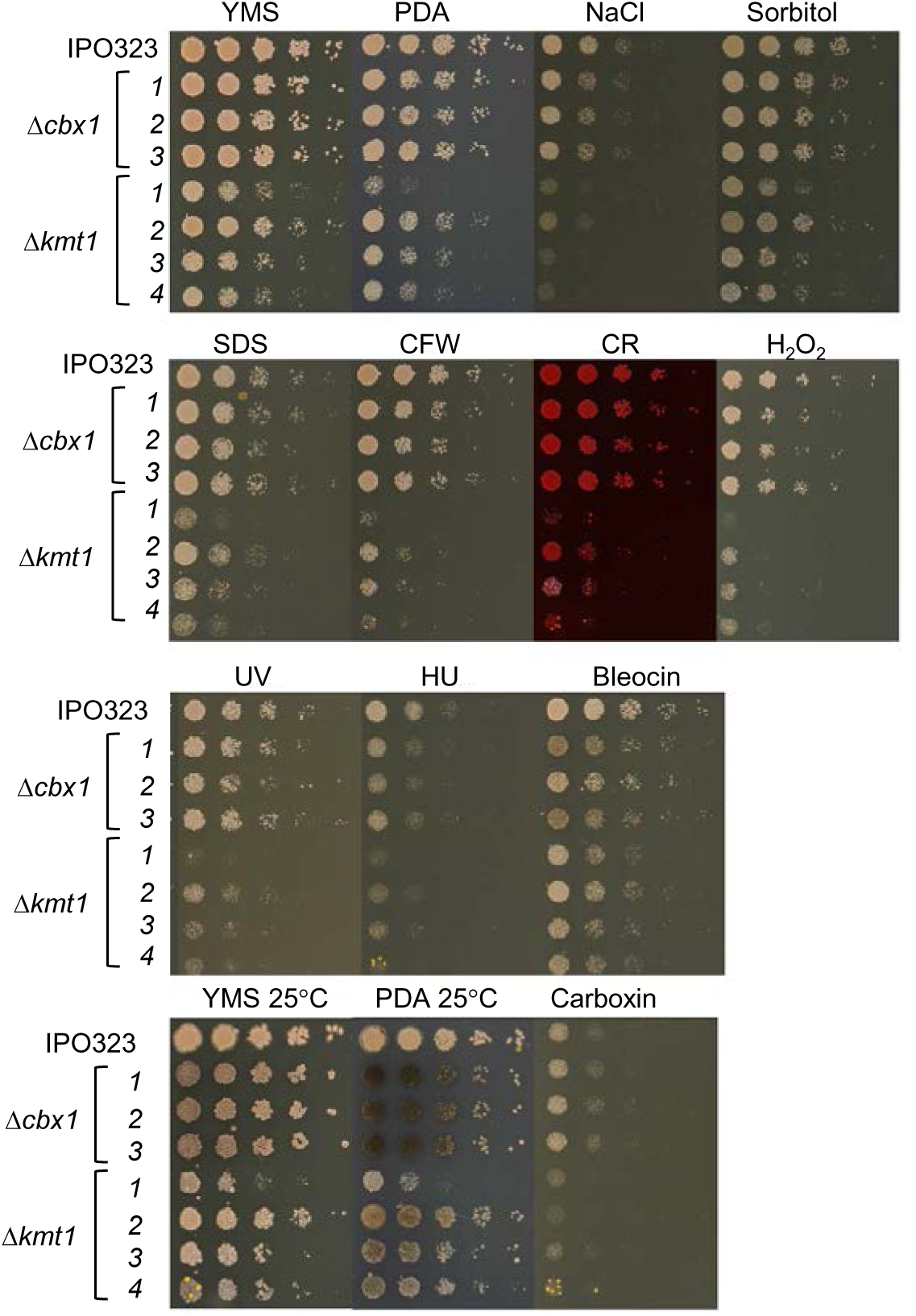
Deletion of *cbx1* and *kmt1* results in distinct phenotypes. Cell suspensions of the indicated strain were subjected to a five-fold serial dilution and pinned onto the indicated agar plates and incubated at 18°C unless indicated otherwise. Agar plates were made with YMS (Yeast extract, malt extract, sucrose) or where indicated, PDA (potato dextrose agar). Concentrations of the stress-inducing agents were, NaCl 1 M, sorbitol 1 M, calcofluor white (CFW) 50 µg/mL, congo red (CR) 150 µg/mL, H_2_O_2_ 2 mM, UV dose 250 J/m^2^, hydroxyurea (HU) 5 mM, Bleocin 250 ng/mL and Carboxin 2.5 ng/mL.

Loss of Kmt1-mediated H3K9 methylation is associated with a severe reduction in virulence (Möller *et al*., 2019) and this was confirmed using our Δ*kmt1* strains (Fig S1B). To determine if Cbx1 is required for the pathogenicity of *Z. triti*ci, wheat infection assays were carried out with the Δ*cbx1* strains. Disease symptoms presented in leaves treated with both the reference (IPO323) and Δ*cbx1* strains, but the onset of symptoms was delayed in the latter (Fig 4A). This was apparent at 14 days post inoculation (dpi) where the areas of leaf covered by necrotic lesions were reduced in the leaves treated with Δ*cbx1* mutants (Fig 4A). Furthermore, at 21 dpi, which typically marks the endpoint of infection, Δ*cbx1* treated leaves had a reduction in the number of visible pycnidia present on the leaf (Fig 4B & C). Therefore, removal of the HP1 homolog Cbx1 results in reduced virulence, but it does not abolish virulence as is the case for the loss of Kmt1. As such the deletion of *cbx1* does not phenocopy the loss of *kmt1* suggesting that H3K9me marks do not exclusively mediate their downstream biological effects through the recruitment of Cbx1.

**Figure 4.**
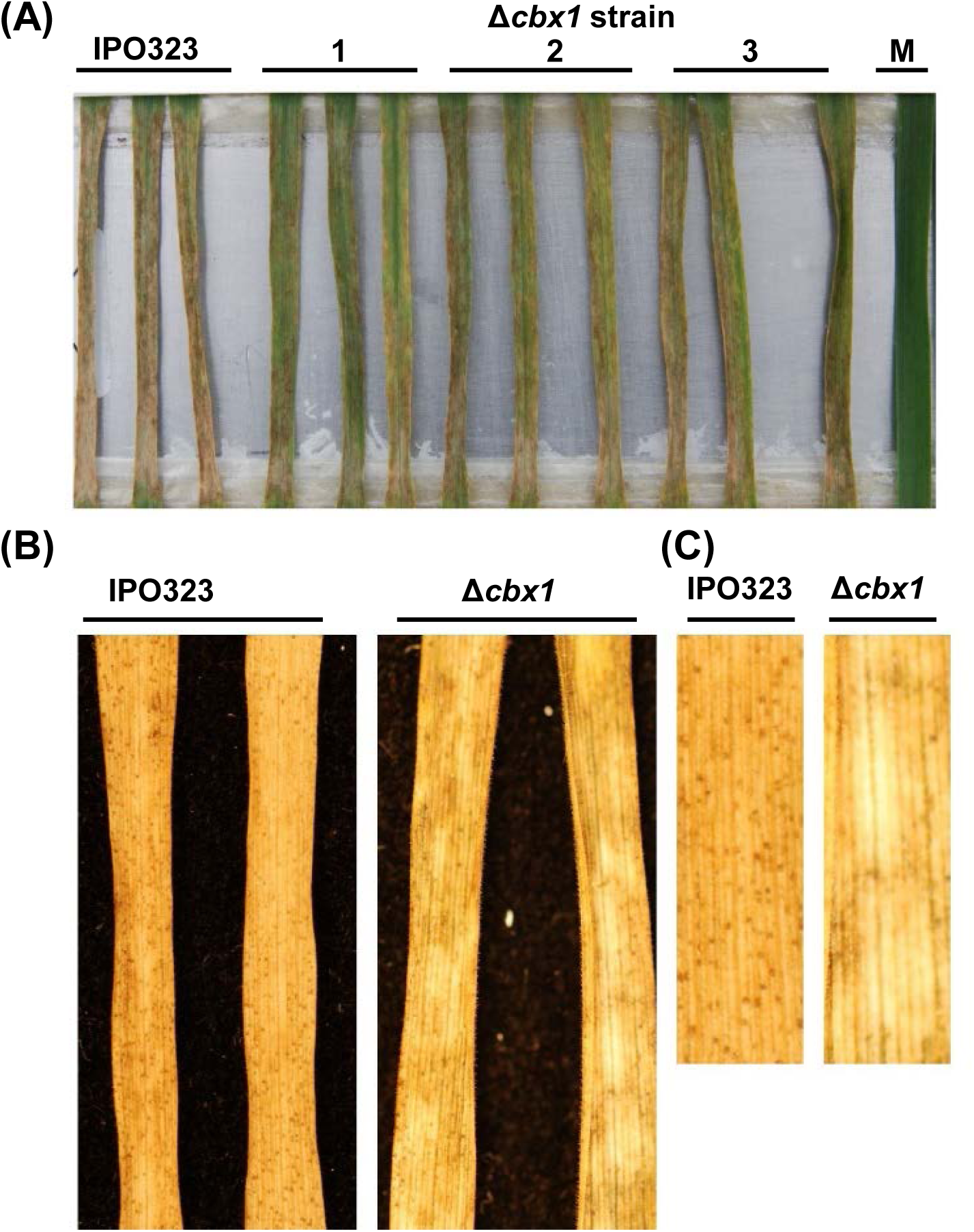
*cbx1* deletion slows disease progression. **(A)** Wheat leaves treated with IPO323, the indicated Δ*cbx1* strains, and a mock infected leaf at 14 days post infection (dpi). **(B)** Wheat leaves from (A) at 21 dpi. A large reduction in pycnidia was observed in leaves treated with the Δ*cbx1* strains. The displayed leaves are representative of three biological repeats. **(C)** Close up of leaves shown in (B).

### Cbx1 and Kmt1 regulate the expression of overlapping sets of genes

To further understand the relationship between Cbx1 and Kmt1, their impacts upon the transcriptome were determined using RNA-seq analysis. RNA was analysed from two biological replicates of two independent isolates of Δ*cbx1* and we also sequenced RNA from two biological replicates of a Δ*kmt1* mutant and the reference IPO323 strain. Principal component analysis revealed a clear grouping of samples from the Δ*cbx1* isolates and the biological replicates for all strains, indicative of low variation (Fig S2). Next, hierarchical clustering was employed to provide an overview of the global similarities between the transcriptomes of the sequenced strains. This revealed that the transcript profiles of Δ*cbx1* and Δ*kmt1* mutants exhibit an overall similarity. Indeed, the Δ*cbx1* mutant profiles were found to be more similar to the Δ*kmt1* strain than to the reference strain (Fig 5A).

**Figure 5.**
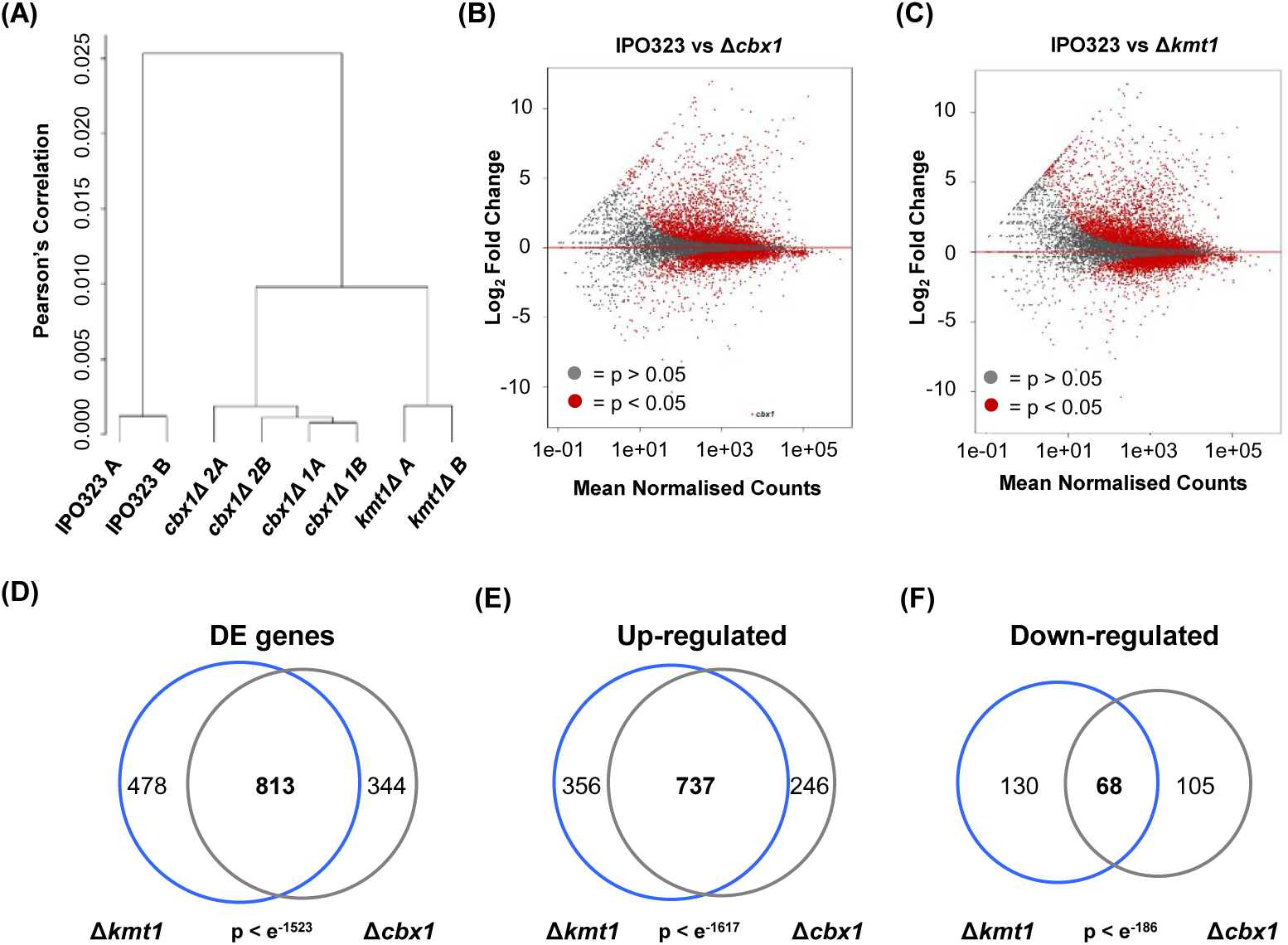
Global impact of Cbx1 and Kmt1 on the transcriptome. **(A)** Hierarchical clustering analysis of the indicated RNA-seq samples. Clustering was performed according to Pearson correlation with the complete linkage method and calculated on log_2_ normalised read counts. **(B)** An MA plot of mean normalised counts plotted against log_2_ fold change for all genes expressed in Δ*cbx1* and IPO323. Genes in red indicate a statistically significant change in gene expression (p<0.05). Genes in grey fall below the cut-off adjusted p value **(C)** An MA plot of Δ*kmt1* and IPO323. Details as for (B). **(D)** Venn diagram comparing differentially expressed genes (>2-fold change in expression, adjusted p value < 0.05) in Δ*cbx1* and Δ*kmt1*. The statistical significance of the overlap was calculated using a Fisher’s test based on hypergeometric distribution. **(E)** As for (D) with up-regulated genes. **(F)** As for (D) with down-regulated genes.

The transcriptomes of Δ*cbx1* and Δ*kmt1* mutant strains were further analysed by identification of differentially expressed (DE) transcripts from protein coding genes with DEseq2 (p < 0.05) (Love *et al*., 2014). Global trends in gene expression were visualised using MA plots (Δ*cbx1* vs IPO322 and Δ*kmt1* vs IPO323) (Fig 5B & C). As expected based on previous analysis (Möller *et al*., 2019), the majority of DE transcripts in Δ*kmt1* were upregulated and the Δ*cbx1* DE transcripts also exhibited a similar trend. Furthermore, for both strains the majority of down-regulated transcripts showed a relatively modest (2-5 fold) decrease in abundance. In comparison, a greater proportion of upregulated transcripts exhibited a more marked (5-10 fold) change in levels. Therefore, like Kmt1, the HP1 protein Cbx1 plays an important role in gene silencing in *Z. tritici*.

Genes that were differentially expressed in the Δ*kmt1* and Δ*cbx1* backgrounds were filtered to select only those that exhibited at least a two-fold change in expression. The total number of DE genes in this category in Δ*cbx1* and Δ*kmt1* was 1157 and 1291 respectively (Tables S1 and S2). Of these genes, 813 were differentially expressed in both strains, an overlap which was found to be highly significant (Fig 5D). The lists of DE genes were further filtered to distinguish between up- and down-regulated genes. Significant overlaps between Δ*cbx1* and Δ*kmt1* gene lists were observed in both categories (Fig 5E & F). Nonetheless, we did identify genes that were differentially expressed in Δ*cbx1* but not Δ*kmt1* and vice versa (Table S3 and S4). The existence of these non-overlapping sets of DE genes may, at least in part, explain the differences in the phenotypes associated with Δ*cbx1* and Δ*kmt1* mutants.

The similarity of the RNA-seq data from the Δ*kmt1* strain generated in this study and the Zt09-Δ*kmt1* strain (Möller *et al*., 2019) was also analysed. For this comparison the more stringent cut-offs (4-fold change in expression, adjusted p value < 0.001) employed by Moller *et al*. were used. Despite the potential for differences in strain background and experimental variability, the overlap in DE genes was highly significant and indicative of a high degree of similarity between the Δ*kmt1* mutant analysed in this study and Zt09-Δ*kmt1* (Fig S3). To determine whether increased stringency affected the relationship between Δ*cbx1* and Δ*kmt1* DE genes, the comparison was repeated using the 4-fold change cut-off. This analysis also revealed a highly significant overlap in the DE gene lists (p < 0.001; Fisher’s test). Overall, these findings indicate that Cbx1 and Kmt1 regulate the expression of similar, albeit non identical, sets of protein-coding genes and are therefore consistent with Cbx1 playing a major role in the function of H3Kme2/3 marks in *Z. tritici*.

### Loss of Cbx1 does not result in a global increase in expression from accessory chromosomes or TEs

It has been demonstrated that deletion of *kmt1* results in a global increase in transcripts derived from the heterochromatin- and TE-enriched accessory chromosomes (Möller *et al*., 2019). To determine whether this was also the case for mutants lacking *cbx1*, normalised read counts were mapped from genes on accessory chromosomes. As expected a significant increase in read counts from accessory chromosomes was observed in the Δ*kmt1* mutant compared to IPO323 (p < 0.014; ANOVA). In contrast no significant increase (p < 0.29; ANOVA) was observed in the Δ*cbx1* background (Fig 6A). Therefore, loss of Cbx1 is not sufficient for a global increase in expression from genes on accessory chromosomes.

**Figure 6.**
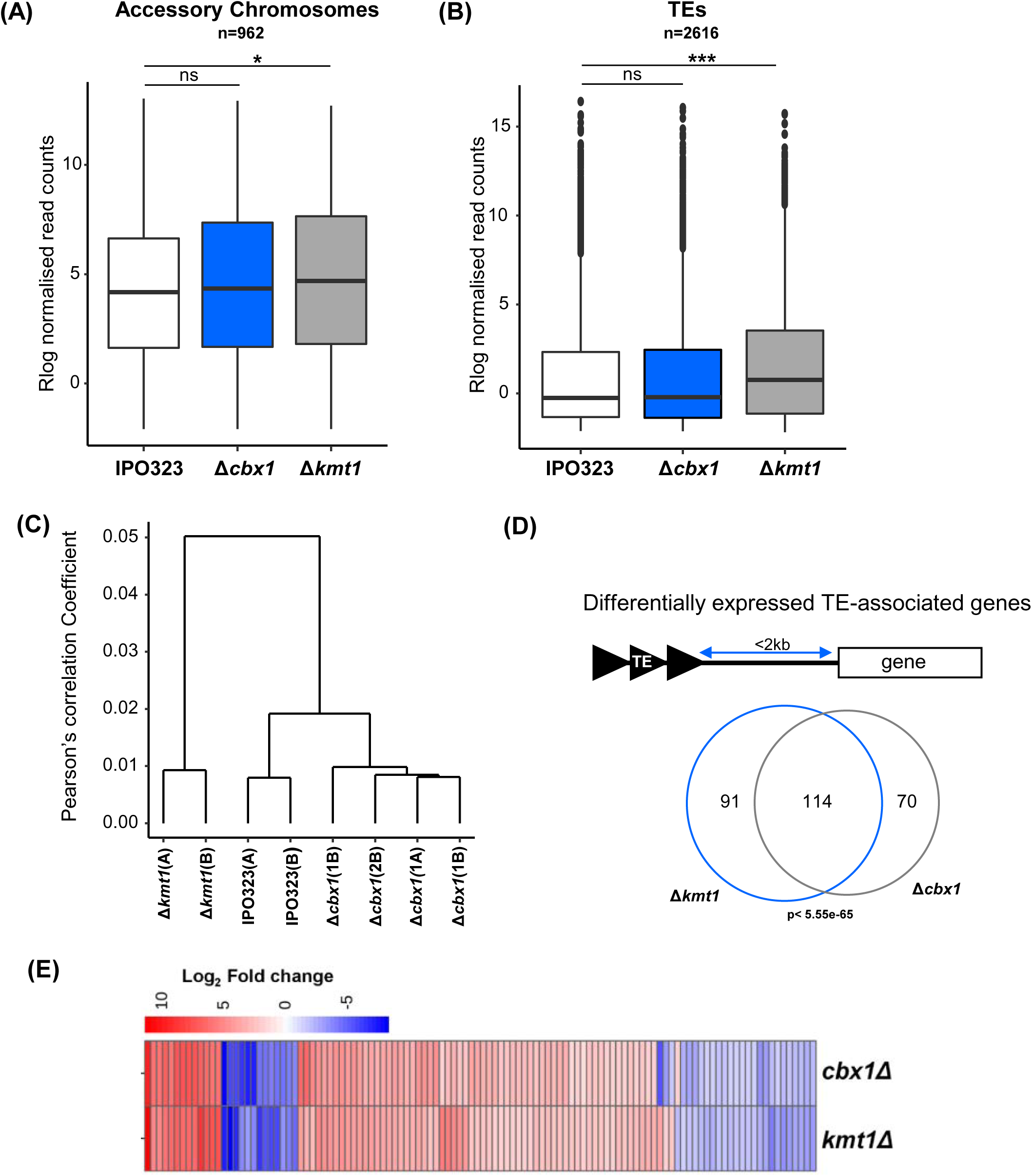
Loss of Cbx1 does not result in a global increase of expression from TEs and accessory chromosomes. **(A)** Median sorted boxplots of Rlog normalised read counts from genes on accessory chromosomes. n = number of genes/elements analysed. * = p < 0.05, ns = not significant (ANOVA). **(B)** TE expression analysed as described in (A) *** = p < 0.001 (ANOVA). **(C)** Hierarchical clustering analysis of TE expression in the indicated RNA-seq samples. Clustering was performed according to Pearson correlation with the complete linkage method and calculated on log_2_ normalised read counts. **(D)** Genes associated (± 2Kb) with TE elements. Venn diagram displaying proportions of TE-associated genes commonly and uniquely differentially expressed in Δ*cbx1* and Δ*kmt1.* The statistical significance of the overlap was calculated using a Fisher test based on hypergeometric distribution. **(E)** Heatmap of TE-associated genes that are differentially expressed in both Δ*cbx1* and Δ*kmt1*. The colour scale represents log_2_ fold changes from -7 to 10. Genes linked to secondary metabolism are indicated. Genes were clustered using euclidean distance and complete linkage.

The accessory chromosomes of *Z. tritici* are highly enriched in TEs and previously it has been shown that expression from these elements is suppressed by H3K9me2/3 (Möller *et al*., 2019). We therefore determined the effect of Cbx1 on the global level of transcripts derived from TEs. As previously observed (Möller *et al*., 2019), a significant net increase in the expression of TEs was detected in Δ*kmt1* (p < 8.25e-09 ANOVA), however in contrast, no significant global increase the Δ*cbx1* mutant was detected (Fig 6B). Indeed, hierarchical clustering revealed that with respect to the profile of TE expression, the Δ*cbx1* mutant is more similar to the reference IPO323 strain than to the Δ*kmt1* mutant (Fig 6C). Therefore, although global silencing of TEs in *Z. tritici* requires Kmt1, and by implication H3K9me2/3, it is not dependent upon recognition of these histone modifications by the HP1 protein, Cbx1.

### Cbx1 regulates the expression of a significant proportion of TE-associated genes

An increased frequency of recombination is often observed around loci surrounding TEs. Genetic instability around such loci in filamentous fungi has been proposed to drive rapid evolution and aid niche adaptation (Dong *et al*., 2015, Faino *et al*., 2016, Laurent *et al*., 2018). Therefore, we analysed the expression of all genes within 2 kb of a TE. In total 1505 genes were identified as ‘TE-associated’ of which 184 were differentially expressed in Δ*cbx1* and 205 in Δ*kmt1*, a highly significant enrichment in both cases (p < 4.35e-07 and p < 9.66e-08 respectively). Furthermore, 114 TE-associated genes were found to be commonly differentially expressed in Δ*cbx1* and Δ*kmt1* (p < 5.55e-65) (Fig 6D). The expression profile of these genes was also found to be highly similar between the Δ*cbx1* and Δ*kmt1* mutants and indeed all but 4 genes exhibited similar expression patterns (Fig 6E). GO-term analysis of the differentially expressed TE–associated genes revealed that 8 of the 114 were annotated as having functions relating to secondary metabolism. However, the majority of these were ‘orphan’ genes with no assigned GO terms. This is not unexpected given that TE-associated and heterochromatic loci are known to be enriched with ‘orphan’ genes in a variety of plant pathogenic fungi (Dong *et al*., 2015).

### Protein encoding genes that are differentially expressed in Δ*cbx1* exhibit only a weak correlation with H3K9me

The removal of HP1 tends to result in the upregulation of genes that are associated with H3K9me2/3-marked chromatin (Chujo & Scott, 2014, Reyes-Dominguez *et al*., 2010, Soyer *et al*., 2014). To investigate whether this is the case in *Z. tritici*, genes that were partially (>1 bp) or completely associated with H3K9me were identified through analysis of published ChIP-seq data (Schotanus *et al*., 2015). The relationship between H3K9me-associated genes and genes that are differentially expressed in Δ*cbx1* was then determined. A total of 247 genes were found to be partially associated with H3K9me (>1 bp) while only 112 were completely associated with this modification. Of the partially H3K9me-associated genes only 31 were differentially expressed in Δ*cbx1*, an overlap which was just statistically significant (p < 0.023) (Fig 7A). The overlap between the completely H3K9me-associated genes and Δ*cbx1* DE genes was also modest (17 genes, p < 0.016) (Fig 7B). At first glance this weak correlation is surprising, however it has previously been observed that deletion of *kmt1* is not sufficient for the upregulation of the majority of H3K9me- associated genes in *Z. tritici* (Möller *et al*., 2019) and analysis of the Δ*kmt1* RNA-seq data generated in this was consistent with these findings. Only a very modest overlap was observed between Δ*kmt1* DE genes and genes that are fully associated with H3K9me and no significant overlap was observed with partially associated genes (Fig 7C & D). Overall, only a very small number of protein coding genes are located in H3K9me-marked chromatin in *Z. tritici* and under *in vitro* growth conditions, the disruption of heterochromatin is insufficient to activate their expression.

**Figure 7.**
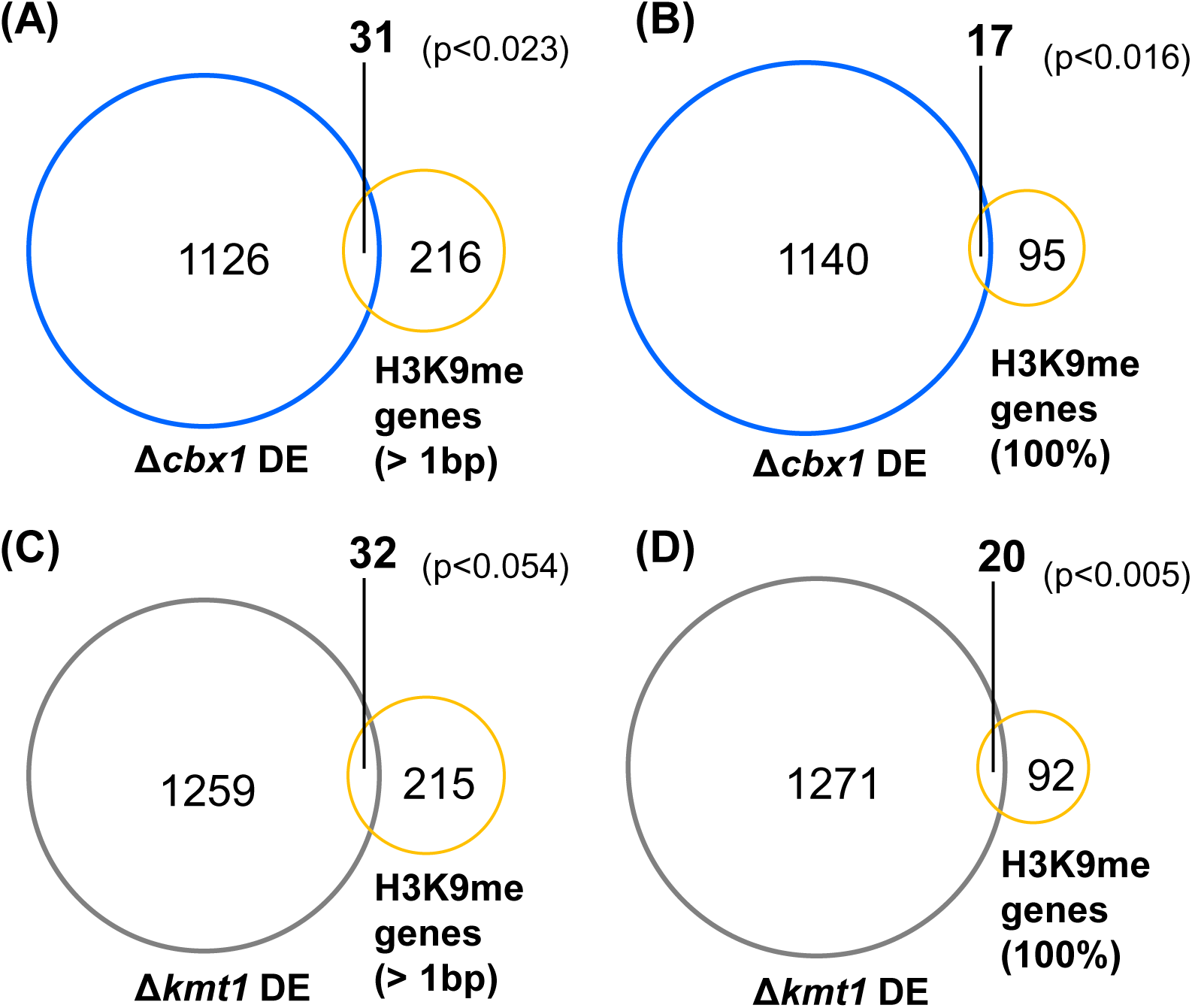
The majority of H3K9me-associated genes are not differentially expressed in Δ*cbx1* mutants. **(A)** Venn diagram of genes differentially expressed (DE) in Δ*cbx1* and genes associated with (> 1bp) H3K9me (B) Δ*cbx1* DE genes and fully marked (100%) H3K9me genes. (C) Δ*kmt1* DE genes and H3K9me3 associated genes. (D) Δ*kmt1* DE genes and H3K9me fully marked genes. The statistical significance of the overlaps in each case was calculated using a Fisher’s test based on hypergeometric distribution.

### Cbx2, a fungal-specific CD protein that binds to H3K9me3

Comparison of the phenotypes of Δ*kmt1* and Δ*cbx1* mutants suggested that some of the downstream effects of H3K9me2/3 histone modifications are likely to be mediated independently of the HP1 homolog Cbx1. One explanation for this would be that *Z. tritci* has additional H3K9me2/3 reader proteins. Therefore, we used BLAST analyses to search for further proteins with the potential to bind H3K9me2/3 PTMs and identified five hypothetical proteins with CD domains (as predicted by ExPASy Prosite and or Pfam). None of these proteins contained a recognisable CSD, consistent with Cbx1 being the sole HP1 isoform in *Z. tritici*. Four of the hypothetical CD proteins were eliminated from further analysis for one or more of the following reasons, (i) they exhibited similarity to retroviral/retrotransposon integrases, (ii) the CD domain lacked critical key aromatic methyl-lysine caging residues or (iii) they were encoded on an accessory chromosome. The remaining hypothetical protein (Mycgr3G108849, hereafter called Cbx2) was predicted to be 703 amino acids in length and have two CD domains in the C-terminal region (Fig 8A and Fig S4). BLAST analyses revealed that organisms that encode proteins with homology to Cbx2 extending beyond the CD domains are limited to species in just a few fungal families (principally the *Mycosphaerellaceae* and *Teratosphaeriaceae*) (Fig 8B and Fig S5). Therefore, unlike the broadly conserved HP1 family member Cbx1, Cbx2 is a fungal-specific CD protein. Sequence analysis revealed that both Cbx2 CDs possess the conserved ‘aromatic cage’ residues that facilitate methyl-lysine binding (Fig S4) and furthermore, chromodomain 1 (CD1) was predicted to be acidic, a characteristic of HP1-type H3K9me binding proteins (Hiragami-Hamada & Nakayama, 2019). Therefore, we investigated the histone binding preferences of Cbx2. A region that encompassed both CD domains (amino acids 503 to 703), was expressed as a GST fusion protein in *E. coli* and purified. Pull-down assays indicated that this domain of Cbx2 binds to histone H3 peptides that are methylated at lysine 9. However, Cbx2 exhibited a clear preference for H3K9me3 relative to H3K9me2 and no specificity for any other tested modification was observed (Fig 8C & D).

**Figure 8.**
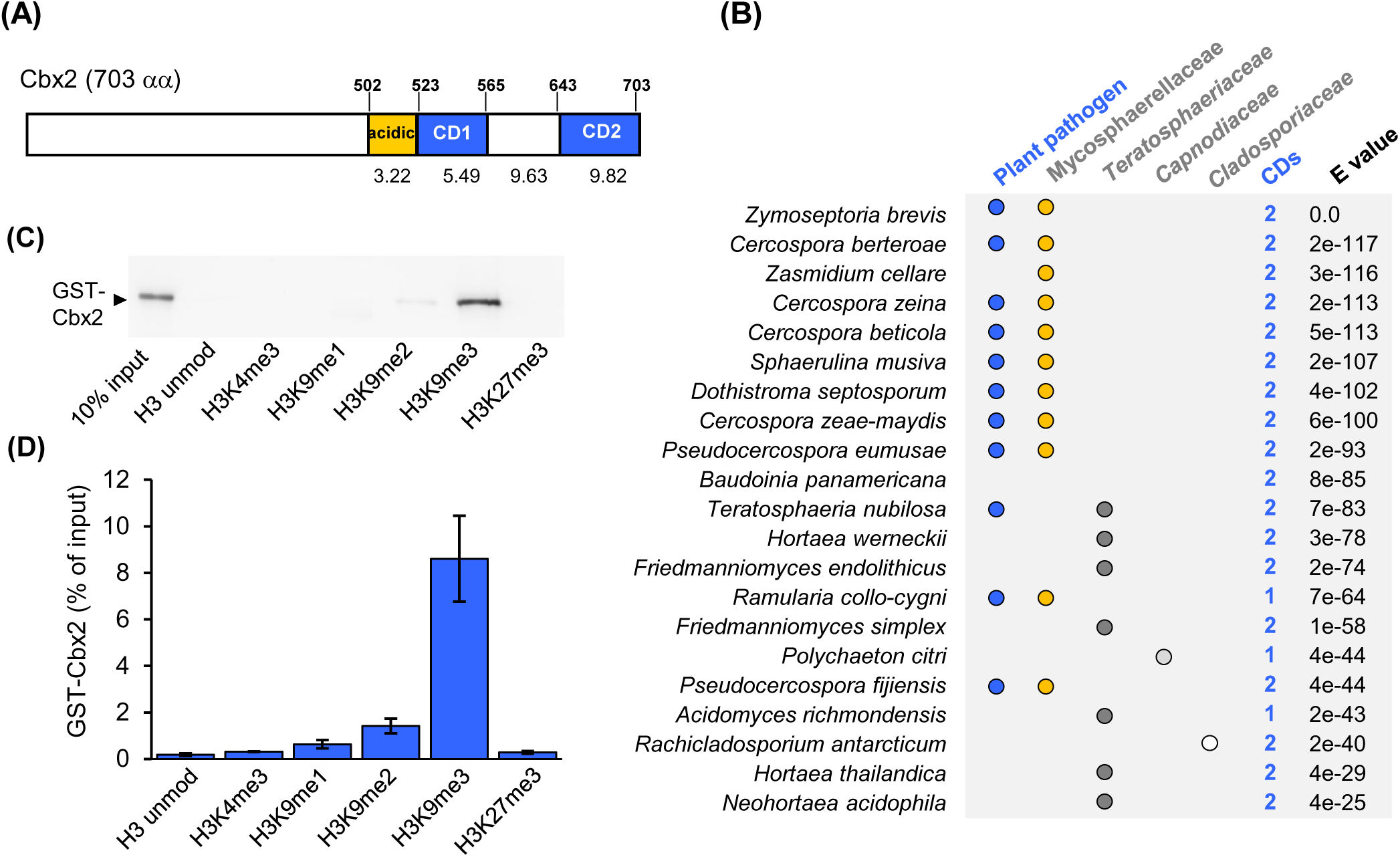
Cbx2, a fungal specific chromodomain protein that binds to H3K9me3 *in vitro*. **(A)** Schematic representation of the domain architecture of Cbx2 showing the location and theoretical pIs of the two chromodomains CD1 and CD2. **(B)** Fungal species with close homologs of Cbx2. The organism, pathogenicity, family, number of chromodomains (predicted by Prosite) and E value relative to Cbx2 are shown. **(C)** GST-Cbx2 (αα 503-703) (1 μg) was incubated with the indicated biotinylated histone H3 peptide and streptavidin beads. Beads were recovered and co-precipitation of GST-Cbx2 was analyzed by western blotting using a GST specific antibody. A 10% input GST-Cbx2 was included as a reference. A representative of three biological repeats is shown. **(D)** Quantification of the GST-Cbx2 signal was relative to the 10% input. Data is the mean of three biological repeats and error bars are ±SEM.

The histone peptide binding assays suggested that Cbx2 has the potential to function as an effector of H3K9me3 PTMs and so Δ*cbx2* strains were generated. Comparison of the Δ*cbx2* mutant with the IPO323 reference strain indicated that loss of Cbx2 does not result in any detectable reduction in fitness or stress resistance (Fig S6). Furthermore, wheat infection assays revealed that, unlike the Δ*kmt1* and Δ*cbx1* strains, Δ*cbx2* strains exhibited no obvious reduction in virulence (Fig 9A and B). Leaves treated with Δ*cbx2* mutants developed disease symptoms at a very similar rate to those treated with the reference IPO323 strain and there was no major difference in the numbers of pycnidia at 21 dpi (Fig 9C). As such, loss of Cbx2 alone does not obviously impact the growth of *Z. tritici* either *in vitro* or *in planta*. This is perhaps not surprising, as when we analysed the expression of *cbx2* using our RNA seq data, we found that this gene was expressed at similar levels to the H3K9 methyltransferase *kmt1*, but at only ∼2.9% of the level of *cbx1*.

**Figure 9.**
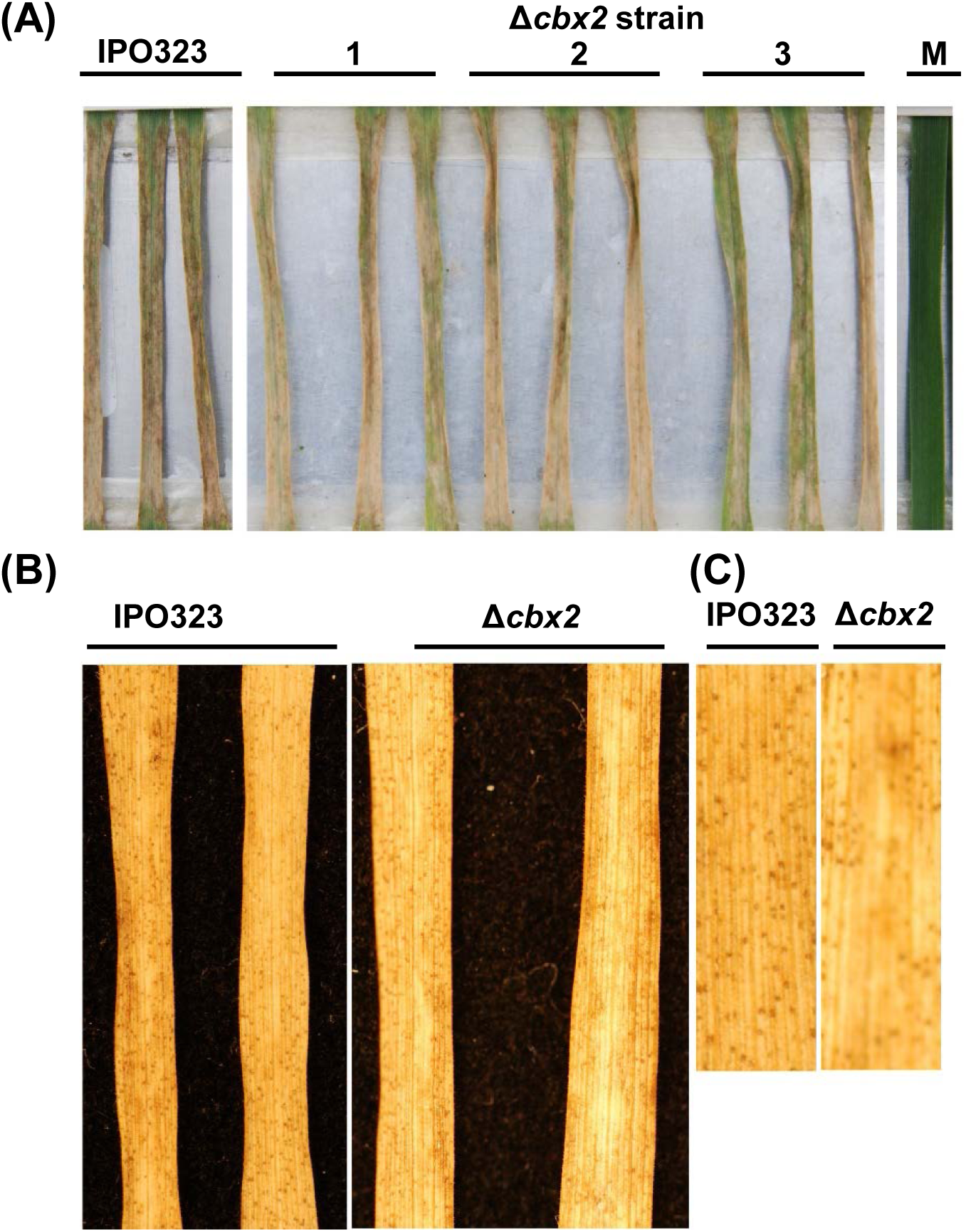
Deletion of *cbx2* does not impair growth *in planta*. **(A)** Wheat leaves treated with IPO323, Δ*cbx2* strains and a mock infection (M) at 14 dpi. **(B)** The same wheat leaves at 21 dpi. The displayed leaves are representative of three biological repeats. **(C)** Close up of the leaves shown in (B).

We hypothesized that Cbx2 may co-operate with Cbx1 but that effects of *cbx2* deletion may be masked when *cbx1* is present. As a test of this, a double deletion mutant was constructed by inserting a *neo* resistance cassette into the *cbx1* locus in the Δ*cbx2* background. Importantly, analysis of the fitness and stress sensitivity profiles of these strains showed that the Δ*cbx1* Δ*cbx2* double mutant has *in vitro* growth phenotypes that closely resemble those associated with Δ*kmt1.* Like the Δ*kmt1* strains, Δ*cbx1* Δ*cbx2* double mutant strains had a slow growth phenotype and were sensitive to osmotic stress (NaCl), oxidative stress (H_2_O_2_), cell wall damaging agents (Calcofluor and Congo Red) and genotoxic agents (HU and Bleomycin) (Fig 10A). As such the deletion of *cbx1* and *cbx2* in combination mimics the loss of H3K9me. These results are consistent with a model whereby key functions of H3K9me PTMs are mediated by a combination of the HP1 homolog Cbx1 and the fungal-specific chromodomain protein, Cbx2.

**Figure 10.**
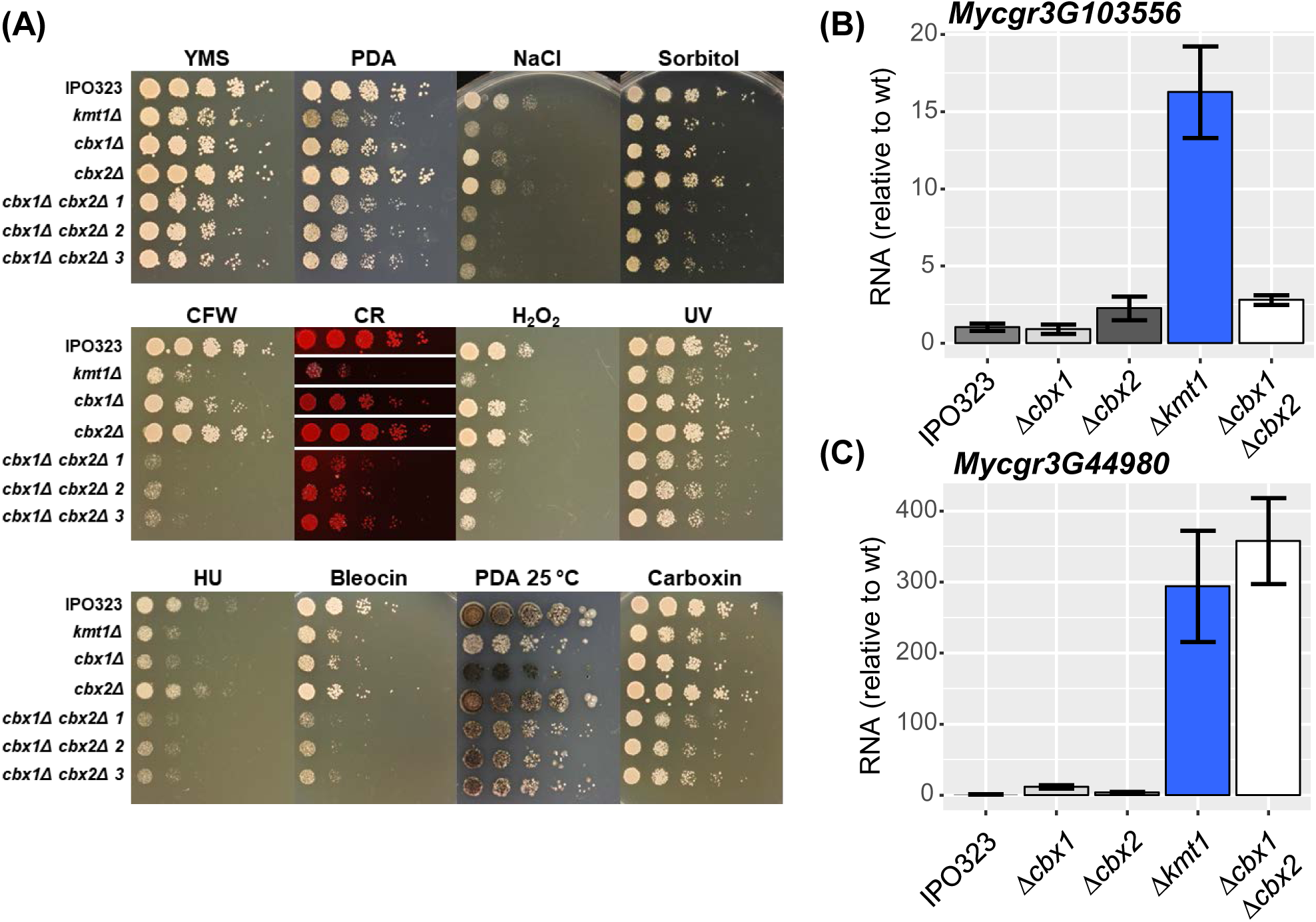
Cbx1 and Cbx2 have redundant functions. **(A)** The *in vitro* growth defects of Δ*cbx1* Δ*cbx2* mutants are similar to Δ*kmt1*. Cell suspensions of the indicated strain were subjected to a five-fold serial dilution and pinned onto the indicated agar plates. Abbreviations and concentrations are as described for Figure 3. **(B)** RNA was extracted from the indicated strains. Mycgr3103556 mRNA levels were determined by RT-qPCR, normalised to actin (Mycgr3G105948) mRNA and scaled relative to the wild type (IPO323) level. Data are the mean of ≥ 3 independent biological repeats and error bars represent ±SEM. **(C)** Mycgr3G44980 mRNA levels were determined as described in (B).

The phenotypes of the Δ*cbx1* Δ*cbx2* double mutant, suggested that Cbx1 and Cbx2 may have redundant functions in gene silencing. We therefore analysed some Kmt1-repressed genes whose expression is not de-repressed by deletion of *cbx1* alone (Fig 5E and Table S4). RT-qPCR analysis showed that the expression of one such Kmt1-repressed gene (*Mycgr3G103556*) was only marginally increased in the Δ*cbx1* Δ*cbx2* double mutant (Fig 10B). In contrast, the deletion of both *cbx1* and *cbx2* genes in combination resulted in an increase in expression of *Mycgr3G44980* comparable to the Δ*kmt1* strain (Fig 10C). Therefore, Cbx1 and Cbx2 do function redundantly to silence the expression of some Kmt1-regulated Z. *tritici* genes.

## DISCUSSION

Heterochromatic H3K9me histone modifications, have a major impact upon the chromosomal stability and virulence of *Z. tritici* (Möller *et al*., 2019). Here we have identified two chromodomain proteins, Cbx1 and Cbx2, which recognize these marks and are implicated in mediating downstream biological events.

Cbx1 bears all the hallmarks of an HP1 ortholog, as it binds to H3K9me2/3 *in vitro* and is enriched at heterochromatic loci. Furthermore, Cbx1 and the H3K9 methyltransferase, Kmt1, regulate the expression of overlapping sets of protein encoding genes. Recognition of H3K9me2/3 modifications by HP1 proteins constitutes a fundamentally conserved step in the formation and function of heterochromatin (Kumar & Kono, 2020). This central role is illustrated by the finding that in some species, such as fission yeast, the phenotypes associated with the loss of HP1 proteins and the respective histone H3K9 methyltransferase are highly similar (Allshire *et al*., 1995). Despite Cbx1 being the sole HP1 homolog in *Z. tritici*, Δ*cbx1* strains have *in vitro* and *in planta* growth defects that are less severe than Δ*kmt1* mutants. These findings are consistent with the data from other plant-associated fungi such as *E. festucae*. While both HepA (HP1) and ClrD (H3K9 methyltransferase) are required for the symbiotic mutualist interaction of *E. festucae* with the grass *Lolium perenne*, Δ*hepA* mutants have only mild defects in axenic culture in comparison to strains lacking Δ*clrD* (Chujo *et al*., 2019). Also RNAi silencing of the HP1 and H3K9 methyltransferase homologs has different effects on the virulence of the oil seed rape pathogen, *L. maculans* (Soyer et al., 2014). It is possible that there are additional H3K9me readers in these organisms.

The transcriptomic analysis revealed a highly significant overlap in the differentially expressed genes in the Δ*kmt1* and Δ*cbx1* backgrounds. Nonetheless, a set of genes was identified whose expression was dependent upon Cbx1, but independent of Kmt1 (Table S3). This suggests that Cbx1 has functions that are independent of H3K9me2/3. Consistent with this, H3K9me-independent roles for HP1 isoforms at telomeres and in DNA damage responses have been reported (Zeng *et al*., 2010). Furthermore, it is well recognised that individual HP1 proteins can be functionally promiscuous and have variety of roles outside of heterochromatin, including transcriptional activation (Zeng *et al*., 2010).

Analysis of the genomic distribution of H3K9me marks in *Z. tritici* has demonstrated that these modifications are predominantly associated with TE elements (Möller *et al*., 2019). Deletion of *kmt1* is associated with a global increase in the abundance of transposon-derived transcripts (Möller *et al*., 2019), a finding that was confirmed in this study. Therefore, in *Z. tritici* as in other eukaryotes, heterochromatin represents a key mechanism for suppressing the activity of repetitive elements. Surprisingly, our findings suggest that Cbx1 is dispensable for the restriction of these elements, at least at a global level and only a small number of TEs are differentially expressed in the Δ*cbx1* strain. *Z. tritici* has up to eight accessory chromosomes that are highly enriched with TEs and are proposed to provide a selective advantage under some environmental conditions (Habig *et al*., 2017). Loss of Kmt1 is associated with a global activation of TEs, elevated loss of accessory chromosomes and wide scale genome rearrangements. The genomic instability in Δ*kmt1* mutants is driven by the redistribution of H3K27me3 modifications which invade regions previously occupied by H3K9me2/3 (Möller *et al*., 2019). That removal of Cbx1 does not result in a global increase in transcripts derived from either TEs or accessory chromosomes, suggests that HP1 function may not be necessary to prevent wide scale re-localization of H3K27me3 modifications. While the loss of H3K9me2/3 severely impacts the ability of *Z. tritici* to colonize wheat leaves, this does not seem to result from mitotic instability as deletion of the H3K27 methyltransferase gene *kmt6* in the Δ*kmt1* background suppresses the elevated level of accessory chromosome loss but does not rescue virulence (Möller *et al*., 2019).

The non-identical phenotypes of the Δ*kmt1* and Δ*cbx1* mutants suggested that additional readers of H3K9me2/3 marks are present in *Z. tritici*. Consistent with this prediction, we have identified Cbx2, a chromodomain protein that recognises H3K9me3 *in vitro*. Unlike Cbx1, which is an HP1 family member and is thus broadly conserved, Cbx2 homologs are restricted to some dothideomycete species suggesting a specialised role in heterochromatin assembly and or maintenance. Also, preliminary evidence suggests that Cbx2 is much less abundant than Cbx1. The *in vitro* binding studies also indicated that the Cbx2 CD region had a preference for H3K9me3 relative to H3K9me2. This is potentially important as analysis of these marks in *S. pombe* has revealed that they demarcate functionally distinct types of heterochromatin that recruit reader proteins with different efficiencies and have different transcriptional silencing potential (Jih *et al*., 2017).

Cbx2 has an unusual structure in that it contains two closely related chromodomains (CD1 and CD2). So far the only characterised proteins that have a double chromodomain structure are the CHD (chromo-ATPase/helicase-DNA-binding) proteins that belong to the SWI/SNF superfamily of ATP-dependent chromatin remodelling enzymes (Yap & Zhou, 2011). It should be noted that the CHD chromodomains belong to a distinct clade that is not involved in the recognition of heterochromatic marks (Yap & Zhou, 2011). The sequences of the Cbx2 CDs are closely related suggesting they arose by duplication. In support of this, some species (e.g. *Polychaeton citri, Ramularia collo-cygni* and *Acidomyces richmondensis*) have Cbx2 homologs that have only a single CD (Fig 8B and Fig S5). It is not yet clear how Cbx2 binds H3K9me3 but it is tempting to suggest that it is achieved via CD1. CD1 is acidic and is flanked by acidic upstream and basic downstream regions, characteristics of H3K9me-binding chromodomains (Hiragami-Hamada & Nakayama, 2019).

While our results suggest that both Cbx1 and Cbx2 are important in executing the functions of H3K9me marks, it is possible that some aspects of their biological function are independent of methyl-lysine reader proteins. Indeed, methylation of lysine 9 may influence transcription or other aspects of chromatin function by preventing the acetylation of this residue. It is also possible that Kmt1 may mediate some functions through the methylation of non-histone targets as has been documented for other SET domain histone methyltransferases (Carlson & Gozani, 2016).

Only a small number of protein encoding genes are located in H3K9me-marked chromatin in *Z. tritici* and consistent with previous findings only a fraction of these genes are differentially expressed in the Δ*kmt1* and Δ*cbx1* mutants. However, analysis of DE genes in Δ*kmt1* has revealed a significant enrichment for genes located in the vicinity of TEs and similar relationship was also observed for Δ*cbx1*. These findings suggest that the heterochromatin associated with TEs can shape the expression of genes in the surrounding chromosomal loci. Indeed, the ability of TE insertions to impact the expression of adjacent genes in *Z. tritici* has been demonstrated (Krishnan *et al*., 2018). It is also noteworthy that *Drosophila* HP1a binds to promoters independently of H3K9me marks and it has been proposed that HP1a then makes transient looping contacts with H3K9me target sites in surrounding regions (Figueiredo *et al*., 2012). This model may explain how H3K9me-marked TEs influence the expression of nearby genes in *Z. tritici*. Furthermore, TEs have been implicated in the organization of loops and other higher order chromosomal structures in a variety of species including fission yeast, flies, plants and mammals (Cam *et al*., 2008, Choudhary *et al*., 2020, Mamillapalli *et al*., 2013, Sun *et al*., 2020).

Analysis of the genomes of fungal phytopathogens has revealed that effector genes tend to be associated with rapidly evolving regions of the genome that are associated with repetitive elements (Dong *et al*., 2015). These observations have led to the suggestion that repetitive elements may organize the regions of genome into functional compartments that drive adaptive evolution. Therefore, it is interesting that the expression of a significant proportion of TE-associated genes is influenced by Kmt1 and Cbx1. Furthermore, recent analysis indicates that genes that are highly expressed at the switch of *Z. tritici* to necrotrophic growth during infection are amongst those most upregulated in the absence of Kmt1 (Soyer *et al*., 2019). Therefore, it will be important to determine how Cbx proteins and the reprogramming of H3K9me- dependent heterochromatin structures contribute to changes in transcriptional programmes during plant infection.

## EXPERIMENTAL PROCEDURES

### Strains, Media and Plasmids

*Zymoseptoria tritici* strains were cultivated on YMS medium (0.4% [w/v] yeast extract, 0.4% [w/v] malt extract, 0.4% [w/v] sucrose) at 18°C in a shaking incubator at 200 rpm. When solid medium was required Bacto agar was added at 2% [w/v]. Gene deletion strains were constructed using *Agrobacterium tumefaciens*-mediated transformations of IPO323 using plasmids derived from pCHYG as previously described (Motteram *et al*., 2009). Flanking regions (>1kb) of the targeted gene were PCR amplified and introduced into pCHYG by Gibson assembly. Gene deletions of *kmt1*, *cbx1* and *cbx2* were constructed by insertion of a hygromycin resistance cassette (*hph*) into the desired locus. The *cbx1 cbx2* double deletion mutant was constructed by inserting a G418 resistance cassette (*neo*) into the *cbx1* locus in the *cbx2* deletion mutant. Correct integration was confirmed by PCR genotyping. Plasmid pCGEN-YR-*cbx1GFP* was constructed by introducing fragments consisting of the *cbx1* promoter, the *cbx1* ORF fused to *EGFP* and 1 kb of terminator sequence from the β-tubulin gene into the *BamHI* site of pC-G418-YR (Sidhu *et al*., 2015) using recombinational cloning in *Saccharomyces cerevisiae*. For the assessment of growth *in planta*, wheat infection assays were performed as previously described (Keon *et al*., 2007).

### *In vitro* sensitivity assays

*Z. tritici* strains were cultured on YMS agar plates for 7 days at 18°C. Cells were then harvested, washed once in sterile 1 x PBS and diluted to OD_600_ of 1.0. Cells were subjected to five-fold serial dilution and pinned onto the indicated YMS and PD agar (Formedium) plates with a 48-pin tool (Sigma). UV irradiation was achieved using a Stratalinker 2400 UV crosslinker (Stratagene). Plates were then incubated for 7 days at 18°C unless otherwise indicated.

### GST fusion proteins

Recombinant Cbx1 was expressed fused to an N-terminal Glutathione-S-Transferase (GST) tag. The *cbx1* sequence (codon optimised for *E. coli*) was synthesized (Eurofins Genomics), cloned into pGEX-6P-1 and transformed in *E. coli* (BL21). Transformants were grown in 2L of LB at 37°C until an OD_600_ of 0.5-0.6 was reached, IPTG was add to a final concentration of 0.5 mM and the culture was incubated at 18°C for 16 hours. The cells were harvested by centrifugation and the resulting pellet was resuspended in 50 mL lysis buffer (50 mM Tris HCl [pH 8.0] 500 mM NaCl 1 mM PMSF), snap frozen in liquid nitrogen and stored at -80. Thawed cell pellets were supplemented with an additional 1 mM PMSF and lysed using a One Shot homogeniser (Constant Systems Ltd) at 20 KPSI at 4°C and then centrifuged at 19 000 RPM in a JA-25.50 rotor (Beckman Coulter) for 30 minutes at 4°C. The supernatant was incubated with 500 µL Pierce^TM^ Glutathione Agarose (Thermo Scientific) pre-equilibrated in wash buffer (50 mM Tris HCl [pH 8.0] 500 mM NaCl) on a rotator for 1 hour. The lysate was then centrifuged at 700 x *g* for 2 minutes and the supernatant discarded. The glutathione agarose was washed once, resuspended in 10 mL wash buffer and applied to a 10 mL disposable gravity flow column (Thermo Scientific). The agarose resin was then washed until a basline A_280_ value was reached. GST fusion protein was eluted off the column in 1 mL fractions using wash buffer supplemented with 10 mM glutathione. Fractions containing GST-Cbx1 were pooled and then subjected to size exclusion chromatography using a Superdex 200 column (GE Lifesciences).

Recombinant GST-Cbx2, composed of an N-terminal GST tag fused to the C-terminal 22 kDa of Cbx2 (amino acids 503 to 703) was produced by Dundee Cell Products.

### Histone H3 peptide binding assays

1 µg of GST tagged protein was added to 1 µg of biotin labelled peptide (EpiCypher) in 300 µL pulldown binding buffer (50 mM Tris [pH 7.5], 300 mM NaCl, 0.01 % NP-40) and rotated overnight at 4°C on a rotating wheel. An aliquot (30 µL) of a 50 % slurry of streptavidin beads (Thermo Scientific) pre-equilibrated in pulldown binding buffer, was added and the sample and incubated at 4°C for one hour on a rotating wheel. Beads were then pelleted by centrifugation at 800 x g for 1 minute and washed four times with 1 mL pulldown binding buffer. The supernatant was removed and streptavidin beads were boiled in 60 µL 2 x protein loading dye (125 mM Tris-HCL [pH 6.8], 20 % glycerol [v/v], 5 % SDS [w/v], 370 mM β-mercaptoethanol [added directly before use]). A control sample of GST fusion protein (100 ng) was loaded alongside the pulldown samples which were resolved on 10% SDS polyacrylamide gels and subject to western blotting using anti-GST antibody (Sigma G7781). Membranes were developed with an ECL plus Chemiluminescent kit (GE Healthcare) and imaged on a Typhoon FLA 9500 (GE Healthcare). GST fusion protein levels relative to the input control were quantified using image J.

### Chromatin Immunoprecipitation (ChIP) Assays

An exponential phase culture of *Z. tritici* was diluted to OD_600_ 0.25, grown overnight at 18°C, and harvested at OD_600_ 0.80. For each 100 mL of culture, 1.35 mL 37 % formaldehyde (Sigma Aldrich) was added to each flask and the cells fixed for 15 minutes. 2 mL 2.5 M glycine was then added to quench the remaining formaldehyde and the flasks incubated at room temperature for a further 5 minutes. Cells were harvested by centrifugation and washed sequentially in 50 mL and 2 mL sterile MilliQ water. Cell pellets were snap-frozen in liquid nitrogen and stored at -80°C. Tissue was ground in under liquid nitrogen in a pestle and mortar and resuspended in freshly made chromatin buffer (50 mM HEPES [pH7.5], 20 mM NaCl, 1 mM EDTA, 1 % Triton X- 100, 0.1 % sodium deoxycholate [w/v]) supplemented with protease inhibitors (1 mM PMSF, 1 μg/mL leupeptin, 1 μg/mL E-64, 0.1 μg/mL pepstatin). CaCl_2_ was then added to a final concentration of 2 mM. To initiate digestion, 150 U MNase (USB/Pharmacia) (prepared as 15 U/μL in 10 mM Tris HCl [pH7.5] 10 mM NaCl 100 μg/ml BSA) per 1 mL of lysate was added and the reaction incubated at 37°C for 20 minutes with frequent mixing by inversion of the tubes. Digestion was stopped by the addition of EGTA to a final concentration of 2 mM. The cell debris was pelleted at 4000 rpm at 4 °C in a microcentrifuge centrifuge and the supernatant was retained. Two 100 μL aliquots of the lysate were taken to be used as ‘input’ and to check MNase digestion respectively. The remaining lysate split was into 200 μL fractions to be used in each immunoprecipitation. To each IP fraction 2 μL of α-GFP antibody (A-11122 – Invitrogen) was added and incubated rotating overnight at 4°C. The following day, 20 μL protein A Dynabeads® (Invitrogen), pre-equilibrated in chromatin buffer were added to each IP and rotated at 4 °C for 2 hours. The supernatant was then removed, and the beads washed twice for 5 minutes at 4 °C in ChIP lysis buffer (50 mM HEPES [pH 7.4], 140 mM NaCl, 1 mM EDTA [8.0], 1 % Triton X-100 [v/v], 0.01 % sodium deoxycholate [w/v]). The beads were then washed once in each of the following buffers: ChIP lysis buffer + 500 mM NaCl, LiCl buffer (10 mM Tris-HCl [pH 8.0], 250 mM LiCl, 0.5% NP40 [v/v], 0.5 % sodium deoxycholate [w/v], 1 mM EDTA), TE buffer (10 mM TRIS [pH 8.0], 1 mM EDTA [pH 8.0]) On the final wash the supernatant was removed and 100 μL 10 % Chelex® 100 (w/v) (Bio-Rad) in MilliQ water was added the beads. 100 μL of 10% Chelex® 100 (w/v) was also added to 10 μL of the input fraction. All samples were then boiled at 100°C for 12 minutes. 2.5 μL 10 mg/mL proteinase K was then added, and the samples incubated at 55°C for 30 minutes. Samples were then boiled at 100°C for 10 minutes, after which the beads and Chelex® were pelleted and 60 μL of the supernatant transferred to a clean tube. The input and IP fractions were diluted by 1:000 and 1:5, respectively. 2 μL of diluted input and IP template were used for each 10 μL qPCR reaction. qPCR was carried out with a KAPA SYBR® FAST qPCR Master Mix Kit with 0.2 mM forward and reverse primers in a Rotor-Gene® 6000 HRM Real Time PCR Machine (Corbett). Primer sequences are detailed in Table S5.

### RNA extraction

Dense liquid cultures of *Z. tritici* were diluted to 0.1 OD_600_ in fresh YMS and grown to OD_600_ 1.0. Cells were then harvested, snap frozen in liquid nitrogen and ground to a fine powder in liquid nitrogen with a pestle and mortar. Approximately 100 mg of ground tissue was added to 2 mL of Tri reagent (Invitrogen), transferred to a 2 mL heavy lock tube (Eppendorf) and centrifuged at 16 000 x g for 5 minutes. The supernatant was extracted with chloroform/isoamyl alcohol and then RNA was precipitated by the addition of an equal volume of propan-2-ol, followed by centrifugation at 16 000 x g for up to 30 minutes. The pellet was washed twice in 70% ethanol, air dried for 15 minutes at room temperature and resuspended in 50 uL of MilliQ water. Aliquots were stored at -80°C until required. For RNA sequencing experiments samples were purified using an RNA Clean and Concentrate column (Zymo research) according to the manufacturer’s instructions.

### RT-qPCR

RNA to be reverse transcribed to cDNA was first treated with Precision DNase (Primer Design) following the manufacturer’s instructions. cDNA was then prepared using SuperScript™ IV Reverse Transcriptase (Invitrogen) with Random Hexamers (Invitrogen) following the manufacturer’s instructions. qPCR was carried out with as described for ChIP assays. Primer sequences are detailed in Table S5.

### RNA-seq and bioinformatics

Purified RNA samples were sequenced by Novogene (China). Read quality was confirmed by FastQC version 0.11.9 (Andrews, 2010) and MultiQC version 1.8 (Ewels *et al*., 2016). STAR version 2.7.1a (Dobin *et al*., 2013) was used to index and align reads to the MG2 IPO323 genome assembly. ‘FeatureCounts’ version 1.6.5 from the Subread package (Liao *et al*., 2014) was used to count reads mapped to genes. Gene annotations (King *et al*., 2017) was used for alignment and read counting. File conversions and manipulations were carried out with SAMtools (Li *et al*., 2009) and BAMtools (Barnett *et al*., 2011). Where necessary BEDTools (Quinlan, 2014) was used to convert between the Zt09 (an IPO323 derivative strain) gene annotation (Grandaubert *et al*., 2015) and the IPO323 strain annotation (King *et al*., 2017). Mapping of reads to transposable elements followed the same analysis pipeline using transposable element annotation (Grandaubert *et al*., 2015). To minimise differences caused by data analysis, published ChIP-seq data (Schotanus *et al*., 2015) was analysed following the previously described workflow (Schotanus *et al*., 2015) with minor modifications including read trimming using Trimmomatic (Bolger *et al*., 2014), alignment with Bowtie 2 version 2.4.1 (Langmead & Salzberg, 2012) and peak coverage determined by RSEG version 0.4.9 (Song & Smith, 2011). Peaks that occurred in both replicates were merged using BEDTools. Gene annotations (King *et al*., 2017) were merged with bed files of identified ChIP-seq peaks to generate lists of genes marked by the specified histone modification (completely or < 1 bp association). Read counts were imported into R, normalised and subject to differential expression analysis with DESeq2 (Love *et al*., 2014). DESeq2 was run independently for protein coding genes and transposable elements. Data manipulation and data plotting were carried out in R with the dplyr, stringr, and ggplot2 packages from the tidyverse (Wickham *et al*., 2019) and the reshape2 package (Wickham, 2007). Heatmaps were made in R with pheatmap (Kolde, 2019). Eggnog mapper (Huerta-Cepas *et al*., 2017) was run to gain additional functional information for the differentially expressed genes.

## DATA AVAILABILITY

The sequence data that support the findings of this study are available at the NCBI Sequence Read Archive (SRA) under accession PRJNA769830. Additional sequence data were derived from resources available in the public domain at the SRA under accessions SRP059394 and PRJNA494102. Other data that support the findings of this study are available from the corresponding author upon reasonable request.

## ACKNOWLEDGEMENTS

We thank Elizabeth Veal for comments on the manuscript. CJF was supported by a BBSRC NLD Doctoral Training Partnership studentship. The authors indicate no conflict of interest.

## AUTHOR CONTRIBUTIONS

CJF and SKW contributed to the conception and design of the study. All authors contributed to the acquisition, analysis, and/or interpretation of the data and writing of the manuscript.

## ABBREVIATED SUMMARY

Heterochromatin associated with methylation of histone H3 on lysine 9 (H3K9me) is required for the genome stability and virulence of the fungal pathogen, *Zymoseptoria tritci.* We have identified chromodomain proteins, Cbx1 and Cbx2, which recognise H3K9me and show that loss of these proteins mimics phenotypes that are associated with the loss of the H3K9 methyltransferase, Kmt1. Overall, our data suggest that key functions of H3K9me modifications are mediated by a combination of Cbx1 and Cbx2.

---

**Figure S1. Deletion of *kmt1* results in loss of H3K9me3 and virulence**

**Figure S2. Principal component analysis (PCA) analysis of RNA-seq data**

**Figure S3. Differentially expressed (DE) genes in Δ*kmt1* overlap with DE genes in the Zt09-Δ*kmt1* strain (Moller *et al*., 2019).**

**Figure S4. Comparison of Cbx2 chromodomains with HP1 chromodomains.**

**Figure S5. Sequence alignments of Cbx2 homologs**

**Figure S6. *In vitro* growth phenotypes of Δ*cbx2* mutants**

**Table S1. Genes differentially expressed in Δ*kmt1***

**Table S2. Genes differentially expressed in Δ*cbx1***

**Table S3. Genes differentially expressed in Δ*cbx1* but not Δ*kmt1***

**Table S4. Genes differentially expressed in Δ*kmt1* but not Δ*cbx1***

**Table S5. Oligonucleotide primers used for qPCR in this study.**

## Notes

### Competing Interest Statement

The authors have declared no competing interest.

## REFERENCES

Allshire, R.C., and Madhani, H.D. (2018) Ten principles of heterochromatin formation and function. Nat Rev Mol Cell Biol 19: 229–244.

Allshire, R.C., Nimmo, E.R., Ekwall, K., Javerzat, J.P., and Cranston, G. (1995) Mutations derepressing silent centromeric domains in fission yeast disrupt chromosome segregation. Genes Dev 9: 218–233.

Andrews, S. (2010) FastQC: a quality control tool for high throughput sequence data Available online at: http://www.bioinformatics.babraham.ac.uk/projects/fastqc.

Bannister, A.J., Zegerman, P., Partridge, J.F., Miska, E.A., Thomas, J.O., Allshire, R.C., and Kouzarides, T. (2001) Selective recognition of methylated lysine 9 on histone H3 by the HP1 chromo domain. Nature 410: 120–124.

Barnett, D.W., Garrison, E.K., Quinlan, A.R., Stromberg, M.P., and Marth, G.T. (2011) BamTools: a C++ API and toolkit for analyzing and managing BAM files. Bioinformatics 27: 1691–1692.

Bolger, A.M., Lohse, M., and Usadel, B. (2014) Trimmomatic: a flexible trimmer for Illumina sequence data. Bioinformatics 30: 2114–2120.

Cam, H.P., Noma, K., Ebina, H., Levin, H.L., and Grewal, S.I. (2008) Host genome surveillance for retrotransposons by transposon-derived proteins. Nature 451: 431–436.

Canzio, D., Larson, A., and Narlikar, G.J. (2014) Mechanisms of functional promiscuity by HP1 proteins. Trends Cell Biol 24: 377–386.

Carlson, S.M., and Gozani, O. (2016) Nonhistone Lysine Methylation in the Regulation of Cancer Pathways. Cold Spring Harb Perspect Med 6.

Choudhary, M.N., Friedman, R.Z., Wang, J.T., Jang, H.S., Zhuo, X., and Wang, T. (2020) Co-opted transposons help perpetuate conserved higher-order chromosomal structures. Genome Biol 21: 16.

Chujo, T., Lukito, Y., Eaton, C.J., Dupont, P.Y., Johnson, L.J., Winter, D., Cox, M.P., and Scott, B. (2019) Complex epigenetic regulation of alkaloid biosynthesis and host interaction by heterochromatin protein I in a fungal endophyte-plant symbiosis. Fungal Genet Biol 125: 71–83.

Chujo, T., and Scott, B. (2014) Histone H3K9 and H3K27 methylation regulates fungal alkaloid biosynthesis in a fungal endophyte-plant symbiosis. Mol Microbiol 92: 413–434.

Cowieson, N.P., Partridge, J.F., Allshire, R.C., and McLaughlin, P.J. (2000) Dimerisation of a chromo shadow domain and distinctions from the chromodomain as revealed by structural analysis. Curr Biol 10: 517–525.

Dobin, A., Davis, C.A., Schlesinger, F., Drenkow, J., Zaleski, C., Jha, S., Batut, P., Chaisson, M., and Gingeras, T.R. (2013) STAR: ultrafast universal RNA-seq aligner. Bioinformatics 29: 15–21.

Dong, S., Raffaele, S., and Kamoun, S. (2015) The two-speed genomes of filamentous pathogens: waltz with plants. Curr Opin Genet Dev 35: 57–65.

Ewels, P., Magnusson, M., Lundin, S., and Kaller, M. (2016) MultiQC: summarize analysis results for multiple tools and samples in a single report. Bioinformatics 32: 3047–3048.

Faino, L., Seidl, M.F., Shi-Kunne, X., Pauper, M., van den Berg, G.C., Wittenberg, A.H., and Thomma, B.P. (2016) Transposons passively and actively contribute to evolution of the two-speed genome of a fungal pathogen. Genome Res 26: 1091–1100.

Figueiredo, M.L., Philip, P., Stenberg, P., and Larsson, J. (2012) HP1a recruitment to promoters is independent of H3K9 methylation in Drosophila melanogaster. PLoS Genet 8: e1003061.

Goodwin, S.B., M’Barek S, B., Dhillon, B., Wittenberg, A.H., Crane, C.F., Hane, J.K., Foster, A.J., Van der Lee, T.A., Grimwood, J., Aerts, A., Antoniw, J., Bailey, A., Bluhm, B., Bowler, J., Bristow, J., van der Burgt, A., Canto-Canche, B., Churchill, A.C., Conde-Ferraez, L., Cools, H.J., Coutinho, P.M., Csukai, M., Dehal, P., De Wit, P., Donzelli, B., van de Geest, H.C., van Ham, R.C., Hammond-Kosack, K.E., Henrissat, B., Kilian, A., Kobayashi, A.K., Koopmann, E., Kourmpetis, Y., Kuzniar, A., Lindquist, E., Lombard, V., Maliepaard, C., Martins, N., Mehrabi, R., Nap, J.P., Ponomarenko, A., Rudd, J.J., Salamov, A., Schmutz, J., Schouten, H.J., Shapiro, H., Stergiopoulos, I., Torriani, S.F., Tu, H., de Vries, R.P., Waalwijk, C., Ware, S.B., Wiebenga, A., Zwiers, L.H., Oliver, R.P., Grigoriev, I.V., and Kema, G.H. (2011) Finished genome of the fungal wheat pathogen Mycosphaerella graminicola reveals dispensome structure, chromosome plasticity, and stealth pathogenesis. PLoS Genet 7: e1002070.

Grandaubert, J., Bhattacharyya, A., and Stukenbrock, E.H. (2015) RNA-seq-Based Gene Annotation and Comparative Genomics of Four Fungal Grass Pathogens in the Genus Zymoseptoria Identify Novel Orphan Genes and Species-Specific Invasions of Transposable Elements. G3 (Bethesda) 5: 1323–1333.

Habig, M., Quade, J., and Stukenbrock, E.H. (2017) Forward Genetics Approach Reveals Host Genotype-Dependent Importance of Accessory Chromosomes in the Fungal Wheat Pathogen Zymoseptoria tritici. mBio 8.

Hiragami-Hamada, K., and Nakayama, J.I. (2019) Do the charges matter?-balancing the charges of the chromodomain proteins on the nucleosome. J Biochem 165: 455–458.

Huerta-Cepas, J., Forslund, K., Coelho, L.P., Szklarczyk, D., Jensen, L.J., von Mering, C., and Bork, P. (2017) Fast Genome-Wide Functional Annotation through Orthology Assignment by eggNOG-Mapper. Mol. Biol. Evol. 34: 2115–2122.

Jih, G., Iglesias, N., Currie, M.A., Bhanu, N.V., Paulo, J.A., Gygi, S.P., Garcia, B.A., and Moazed, D. (2017) Unique roles for histone H3K9me states in RNAi and heritable silencing of transcription. Nature 547: 463–467.

Kellner, R., Bhattacharyya, A., Poppe, S., Hsu, T.Y., Brem, R.B., and Stukenbrock, E.H. (2014) Expression profiling of the wheat pathogen Zymoseptoria tritici reveals genomic patterns of transcription and host-specific regulatory programs. Genome Biol Evol 6: 1353–1365.

Keon, J., Antoniw, J., Carzaniga, R., Deller, S., Ward, J.L., Baker, J.M., Beale, M.H., Hammond-Kosack, K., and Rudd, J.J. (2007) Transcriptional adaptation of Mycosphaerella graminicola to programmed cell death (PCD) of its susceptible wheat host. Mol Plant Microbe Interact 20: 178–193.

King, R., Urban, M., Lauder, R.P., Hawkins, N., Evans, M., Plummer, A., Halsey, K., Lovegrove, A., Hammond-Kosack, K., and Rudd, J.J. (2017) A conserved fungal glycosyltransferase facilitates pathogenesis of plants by enabling hyphal growth on solid surfaces. PLoS Pathog 13: e1006672.

Kolde, R., (2019) pheatmap: Pretty Heatmaps. In., pp.

Krishnan, P., Meile, L., Plissonneau, C., Ma, X., Hartmann, F.E., Croll, D., McDonald, B.A., and Sanchez-Vallet, A. (2018) Transposable element insertions shape gene regulation and melanin production in a fungal pathogen of wheat. BMC Biol 16: 78.

Kumar, A., and Kono, H. (2020) Heterochromatin protein 1 (HP1): interactions with itself and chromatin components. Biophys Rev 12: 387–400.

Langmead, B., and Salzberg, S.L. (2012) Fast gapped-read alignment with Bowtie 2. Nat. Methods 9: 357–359.

Laurent, B., Palaiokostas, C., Spataro, C., Moinard, M., Zehraoui, E., Houston, R.D., and Foulongne-Oriol, M. (2018) High-resolution mapping of the recombination landscape of the phytopathogen Fusarium graminearum suggests two-speed genome evolution. Mol Plant Pathol 19: 341–354.

Lee, W.S., Rudd, J.J., Hammond-Kosack, K.E., and Kanyuka, K. (2014) Mycosphaerella graminicola LysM effector-mediated stealth pathogenesis subverts recognition through both CERK1 and CEBiP homologues in wheat. Mol Plant Microbe Interact 27: 236–243.

Li, H., Handsaker, B., Wysoker, A., Fennell, T., Ruan, J., Homer, N., Marth, G., Abecasis, G., Durbin, R., and Genome Project Data Processing, S. (2009) The Sequence Alignment/Map format and SAMtools. Bioinformatics 25: 2078–2079.

Liao, Y., Smyth, G.K., and Shi, W. (2014) featureCounts: an efficient general purpose program for assigning sequence reads to genomic features. Bioinformatics 30: 923–930.

Love, M.I., Huber, W., and Anders, S. (2014) Moderated estimation of fold change and dispersion for RNA-seq data with DESeq2. Genome Biol 15: 550.

Machida, S., Takizawa, Y., Ishimaru, M., Sugita, Y., Sekine, S., Nakayama, J.I., Wolf, M., and Kurumizaka, H. (2018) Structural Basis of Heterochromatin Formation by Human HP1. Mol Cell 69: 385–397 e388.

Mamillapalli, A., Pathak, R.U., Garapati, H.S., and Mishra, R.K. (2013) Transposable element ’roo’ attaches to nuclear matrix of the Drosophila melanogaster. J Insect Sci 13: 111.

Marshall, R., Kombrink, A., Motteram, J., Loza-Reyes, E., Lucas, J., Hammond- Kosack, K.E., Thomma, B.P., and Rudd, J.J. (2011) Analysis of two in planta expressed LysM effector homologs from the fungus Mycosphaerella graminicola reveals novel functional properties and varying contributions to virulence on wheat. Plant Physiol 156: 756–769.

Möller, M., Schotanus, K., Soyer, J.L., Haueisen, J., Happ, K., Stralucke, M., Happel, P., Smith, K.M., Connolly, L.R., Freitag, M., and Stukenbrock, E.H. (2019) Destabilization of chromosome structure by histone H3 lysine 27 methylation. PLoS Genet 15: e1008093.

Motteram, J., Kufner, I., Deller, S., Brunner, F., Hammond-Kosack, K.E., Nurnberger, T., and Rudd, J.J. (2009) Molecular characterization and functional analysis of MgNLP, the sole NPP1 domain-containing protein, from the fungal wheat leaf pathogen Mycosphaerella graminicola. Mol Plant Microbe Interact 22: 790–799.

Quinlan, A.R. (2014) BEDTools: The Swiss-Army Tool for Genome Feature Analysis. Curr Protoc Bioinformatics 47: 11 12 11–34.

Reyes-Dominguez, Y., Bok, J.W., Berger, H., Shwab, E.K., Basheer, A., Gallmetzer, A., Scazzocchio, C., Keller, N., and Strauss, J. (2010) Heterochromatic marks are associated with the repression of secondary metabolism clusters in Aspergillus nidulans. Mol Microbiol 76: 1376–1386.

Rudd, J.J. (2015) Previous bottlenecks and future solutions to dissecting the Zymoseptoria tritici-wheat host-pathogen interaction. Fungal Genet Biol 79: 24–28.

Rudd, J.J., Kanyuka, K., Hassani-Pak, K., Derbyshire, M., Andongabo, A., Devonshire, J., Lysenko, A., Saqi, M., Desai, N.M., Powers, S.J., Hooper, J., Ambroso, L., Bharti, A., Farmer, A., Hammond-Kosack, K.E., Dietrich, R.A., and Courbot, M. (2015) Transcriptome and metabolite profiling of the infection cycle of Zymoseptoria tritici on wheat reveals a biphasic interaction with plant immunity involving differential pathogen chromosomal contributions and a variation on the hemibiotrophic lifestyle definition. Plant Physiol 167: 1158–1185.

Schotanus, K., Soyer, J.L., Connolly, L.R., Grandaubert, J., Happel, P., Smith, K.M., Freitag, M., and Stukenbrock, E.H. (2015) Histone modifications rather than the novel regional centromeres of Zymoseptoria tritici distinguish core and accessory chromosomes. Epigenetics Chromatin 8: 41.

Sidhu, Y.S., Cairns, T.C., Chaudhari, Y.K., Usher, J., Talbot, N.J., Studholme, D.J., Csukai, M., and Haynes, K. (2015) Exploitation of sulfonylurea resistance marker and non-homologous end joining mutants for functional analysis in Zymoseptoria tritici. Fungal Genet Biol 79: 102–109.

Smothers, J.F., and Henikoff, S. (2000) The HP1 chromo shadow domain binds a consensus peptide pentamer. Curr Biol 10: 27–30.

Song, Q., and Smith, A.D. (2011) Identifying dispersed epigenomic domains from ChIP-Seq data. Bioinformatics 27: 870–871.

Soyer, J.L., El Ghalid, M., Glaser, N., Ollivier, B., Linglin, J., Grandaubert, J., Balesdent, M.H., Connolly, L.R., Freitag, M., Rouxel, T., and Fudal, I. (2014) Epigenetic control of effector gene expression in the plant pathogenic fungus Leptosphaeria maculans. PLoS Genet 10: e1004227.

Soyer, J.L., Grandaubert, J., Haueisen, J., Schotanus, K., and Stukenbrock, E.H. (2019) In planta chromatin immunoprecipitation in Zymoseptoria tritici reveals chromatin-based regulation of putative effector gene expression. BioRxiv.

Soyer, J.L., Rouxel, T., and Fudal, I. (2015) Chromatin-based control of effector gene expression in plant-associated fungi. Curr Opin Plant Biol 26: 51–56.

Steinberg, G. (2015) Cell biology of Zymoseptoria tritici: Pathogen cell organization and wheat infection. Fungal Genet Biol 79: 17–23.

Sun, L., Jing, Y., Liu, X., Li, Q., Xue, Z., Cheng, Z., Wang, D., He, H., and Qian, W. (2020) Heat stress-induced transposon activation correlates with 3D chromatin organization rearrangement in Arabidopsis. Nat Commun 11: 1886.

Turck, F., Roudier, F., Farrona, S., Martin-Magniette, M.L., Guillaume, E., Buisine, N., Gagnot, S., Martienssen, R.A., Coupland, G., and Colot, V. (2007) Arabidopsis TFL2/LHP1 specifically associates with genes marked by trimethylation of histone H3 lysine 27. PLoS Genet 3: e86.

Uhse, S., and Djamei, A. (2018) Effectors of plant-colonizing fungi and beyond. PLoS Pathog 14: e1006992.

Wickham, H. (2007) Reshaping Data with the reshape Package. Journal of Statistical Software 21: 1–20.

Wickham, H., Averick, M., Bryan, J., Chang, W., D’Agostino McGowan, L., François, R., Grolemund, G., Hayes, A., Henry, L., Hester, J., Kuhn, M., Pedersen, T., Miller, E., Bache, S., Müller, K., Ooms, J., Robinson, D., Seidel, P., Spinu, V., Takahashi, K., Vaughan, D., Wilke, C., Woo, K., and Yutani, H. (2019) Welcome to the tidyverse. Journal of Open Source Software 4: 1686.

Yale, K., Tackett, A.J., Neuman, M., Bulley, E., Chait, B.T., and Wiley, E. (2016) Phosphorylation-Dependent Targeting of Tetrahymena HP1 to Condensed Chromatin. mSphere 1.

Yap, K.L., and Zhou, M.M. (2011) Structure and mechanisms of lysine methylation recognition by the chromodomain in gene transcription. Biochemistry 50: 1966–1980.

Zeng, W., Ball, A.R., Jr., and Yokomori, K. (2010) HP1: heterochromatin binding proteins working the genome. Epigenetics 5: 287–292.

