## Supplementary Figures for "The chromodomain proteins, Cbx1 and Cbx2 have distinct roles in the regulation of heterochromatin and virulence in the fungal wheat pathogen, *Zymoseptoria tritici*"

**(A)**

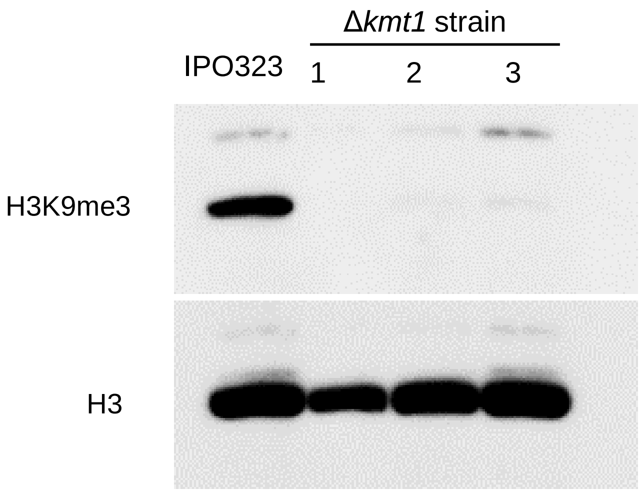


**(B)**

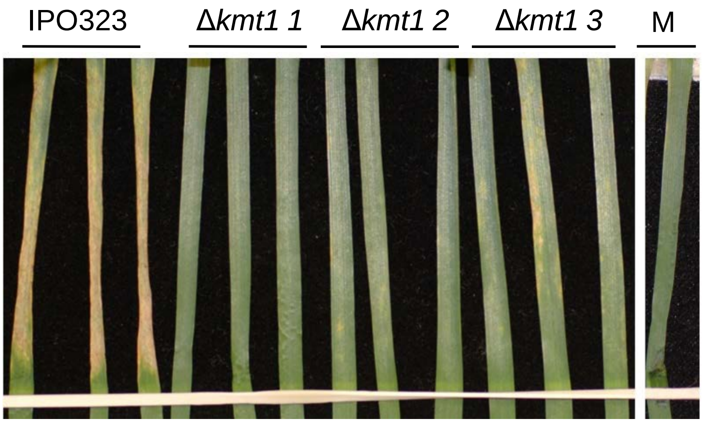


**Figure S1. Deletion of *kmt1* results in loss of H3K9me3 and virulence. (A)** Acid extracted protein samples were prepared from the indicated strains and analysed by immunoblotting with anti-H3K9me3 (Active Motif: 39162) antibodies (upper panel). Total histone H3 levels were determined using antibody specific to the histone H3 C-terminal region (lower panel). **(B)** *In planta* infection assays of Δ*kmt1*. A marked reduction of virulence was observed in wheat leaves treated with Δ*kmt1* cells at 14 days post infection (dpi) relative to the reference IPO323 strain. No further disease progression in leaves treated with Δ*kmt1* strains was observed after 21 dpi (data not shown). Each leaf is representative of a biological repeat (n = 3). M = mock/negative control.

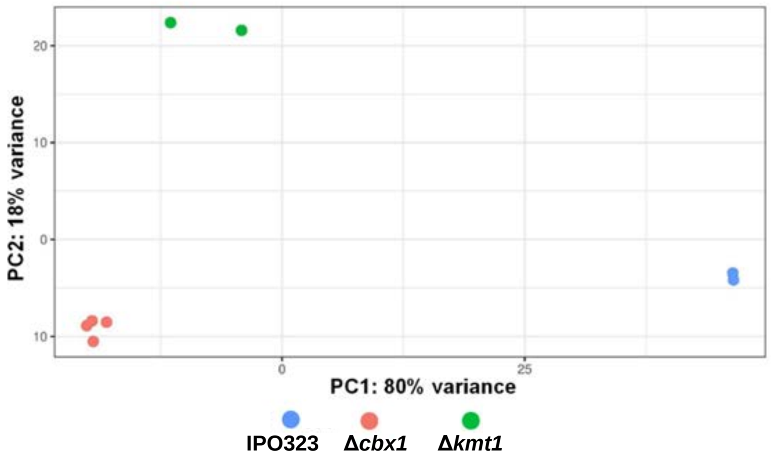


**Figure S2. Principal component analysis (PCA) of RNA-seq data**. Each point represents an RNA-seq sample (as indicated in the key). Clustering of samples is indicative of low variation between replicates and between the individual Δ*cbx1* isolates.

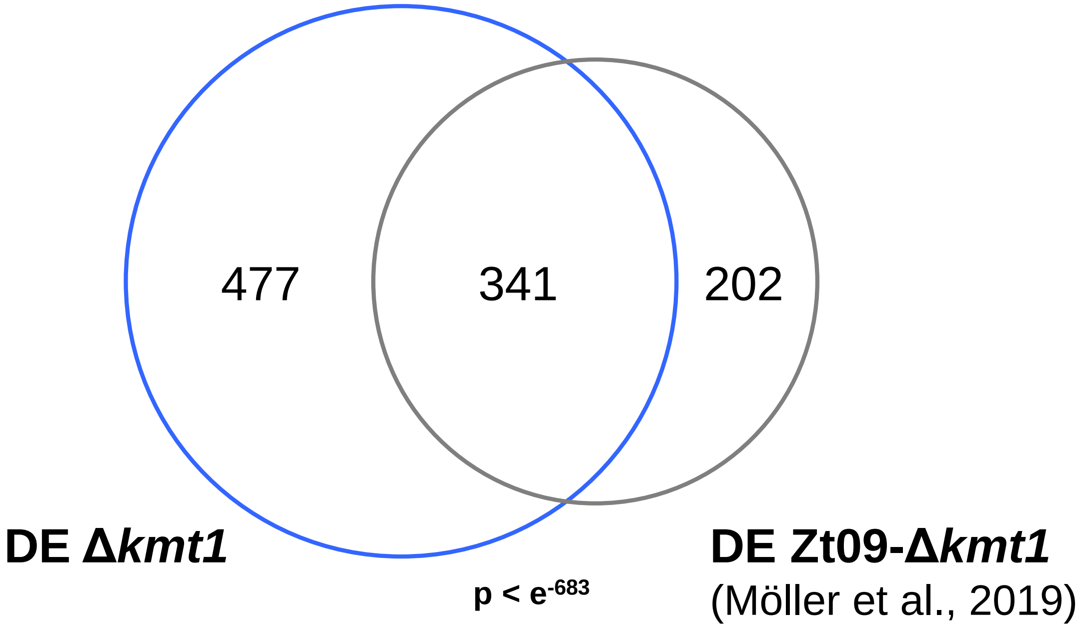


**Figure S3. Differentially expressed (DE) genes in Δ*kmt1* overlap with DE genes in the Zt09-Δ*kmt1* strain (Möller *et al.*, 2019).** DE genes were filtered to match the cut-off criteria from Möller *et al.* (2019) where an expression fold change of 4 and adjusted p-value of 0.001 was applied. The statistical significance of the overlap was calculated using a Fisher's test based on hypergeometric distribution.

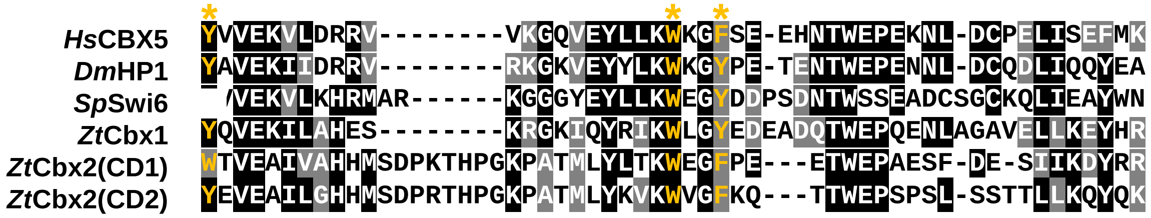


**Figure S4. Comparison of Cbx2 chromodomains with HP1 chromodomains.** Sequence alignment of the chromodomains from the indicated proteins was generated using CLUSTAL. Full shading (black) represents conservation of an amino acid in at least 50% of the sequences whilst grey shading denotes conservation of a residue of similar chemistry in at least 50% of the analysed sequences. Aromatic 'methyl-lysine cage' residues are coloured yellow and their position is highlighted with an asterisk.

```
H.werneckii        1 ----------------------------------------------------MSILASVNVPMYS-PA--
T.nubilosa         1 ----------------------------------------------------MPPLAVVEIPVLS-PS--
B.panamericana     1 ----------------------------------------------------MPPLAHVKIPLYI-PT--
F.endolithicus     1 ----------------------------------------------------MPTLARVYVPSYD-RS--
F.simplex          1 ------------------------------------GLAQLADSTNGTWASKMPTLATVYVPL---RS--
R.collo-cygni      1 ----------------------------------------------------------------------
Z.tritici          1 ----------------------------------------------------MPPALPDTGIIDLFTSPE
Z.brevis           1 ----------------------------------------------------MPPALPDSGIIDLSTSPE
Zas.cellare        1 ----------------------------------------------------MPPLLTSDIEKP------
D.septosporum      1 ----------------------------------------------------MPPITGLTIP--------
P.eumusae         71 AHPRLHFIKRTRFLTHTHTHTYTHTCTYTHFLLLPPALRCICRELLMSWL-NMPPSLESSLALNDPS---
S.musiva           1 ----------------------------------------------------MPPSLEVTQARNKTP---
C.berteroae        1 ----------------------------------------------------MPPSLDATLYRERTP---
C.beticola         1 ----------------------------------------------------MPPSLDATLYREKTP---
C.zeina            1 ----------------------------------------------------MPPSLDATLLRESMA---
C.zeae-maydis      1 ----------------------------------------------------MPPSLDSILFREKIP---

H.werneckii       16 ---------------TS---------KGKAAARASP-PRRVYGDEPLPVKLSDVAPRG---AQIVGRVLG
T.nubilosa        16 ---------------PASSTPPPKQDKGKATVKRQPERRDKENSKPLSIRLDQDAPRA---ATIIAREIT
B.panamericana    16 ---------------LPVSRQSHAGTS--YESKAQY-RPRVYGDAPLSVKPSEDAPAN---ARIIGRVIE
F.endolithicus    16 ---------------PSTATDWSSQSR--VSRRSLP-MPRTYGKGPLSVKPSENSSVN---ARIVGRVVG
F.simplex         30 ---------------PSTATSSSGNSR--VSVKPQP-VLRTYGKEPLPVKPSADSSVN---ARIVGRVLG
R.collo-cygni      1 ----------------------------------------------------------------------
Z.tritici         19 QSPDPPPARK--ALFPVIDRSPSR--------KVPW-IPRTYGDEPLPVRLSTQAPRNAKDARIVGRVE-
Z.brevis          19 QSPEPPPARK--ALFPVIDRSPAR--------KVPW-VPRTYGDEPLPIRLSTQAPRNAKDARIVGRVNA
Zas.cellare       13 VDR------SPHRMVPVIKVPPHP-----PVSKPAP-QPRTYSDEPLPVRPSDHAPRK---AAIKERIAL
D.septosporum     11 VYR-----------ASESPECPRQSLSA-KSVKLRN-QPRIYQDGPLPIKLSDHAPRQ---ARITGRVAT
P.eumusae        137 TIK----RKQLVLDVPLIRKSPIKSI------GKWP-KPRMYGDEPLPVKPSDVNPRN---GRIVGRVAE
S.musiva          16 VFKQPLVQRRLVFDDAVVERSPQRTISNNHKGKRAV-KPRTYGEGPLPFKPSDTAPRN---ARIVGRVAE
C.berteroae       16 VFKHPPLQKRVVLGSGVISRSPERGTNAK--SKNAL-KPRSYGDGPLPFKPSETAPRD---ARIVGRVAE
C.beticola        16 VFKHSPLQKRVVLGSGVISRSPERGTNAK--SKNAL-KPRSYGDGPLPFKPSETAPRD---ARIVGRVAE
C.zeina           16 VLKQPPLQKKLVLGKDVISRSPERGSTAQTKIKSAL-KPRRYGDGPLPSKPSETAPRN---AQIVGRVAE
C.zeae-maydis     16 VSKQHSLQRRLVLGDSVISRSPERGIAAKTKTNGAP-KPRSYGDGPLPFKPSETAPRN---AHIVGRVAE

H.werneckii       58 SS--G-AFYTLKIGDTTLEDVAVDEVLDYISADHLEEYETHQFAEEAELLRIAETENERMERERQEMKRE
T.nubilosa        68 DR--G-ALYTLRIGEVEVPQVELDEILHYVSPEHLELYESQQFIEEEEAQRVAEEAEMQMRLAKLERMKQ
B.panamericana    65 PT--G-SVYQLRIGDVEIHDVGIDEILDYVSPECLEEYEHQQFEEEAELRRIAEIEAERQEAERRERQRH
F.endolithicus    65 DH--G-GRYSLKIGNEALENVGVEEILDYVSADHLEDYENQQFEEEREVRRVAKE---ILAYEKMQRRNA
F.simplex         79 DR--G-GFYSLKIGNEELNNVGVEEILDYVSADHLEDYENQQFAEETEVKRVAKLNEERLAIEKQERRAE
R.collo-cygni      1 -----------MVGQSQITDVSVTEILQFVSPYDLEEFENGQFLEEEENRKIAQAAEEAAEEARRLRKLE
Z.tritici         77 --DGRKVTYTVMLGTTEVSEVDLDEILTYVSPYDLEEYEHEQFKVDGEKRRIDEEAKQAERERKLERLKE
Z.brevis          78 TGEGGKVTYTVMLGSTEVSEVDLDEILTYVSPYDLEEYENEQFKVDGENRRAVEVAKQAERERKLERQKD
Zas.cellare       68 PD--RV-AYTLDIGGVEIEDVGIHEVLDYVSAYDLETFEHRQFEEEREIMRIANLEQEAE----RERRKQ
D.septosporum     65 PD--KV-AYTIEIGGVIMNDVGIDEILDYVSPFELERFETQQFVEEIEIDRIAQEAADAEAEHKREQQKI
P.eumusae        193 FG--QIPVYTLAVGDTEINGIKLWEILDYVSPLELERFENQQFKEEAEALEIARAAAEIEKERKRLKQAE
S.musiva          82 PG--KQPIYSLKVGDIQVDDVCLHDILEYVSPYELERYEHQQFEEEREVLNASLAAAAEEEEKRRERHKG
C.berteroae       80 PG--KLPMFTLKVGDVEMEDITLHEILEYVSPYELERFENEQFAEEREALAVALAAAEAEDERRRQRRKE
C.beticola        80 PG--KLPMFTLKVGDVEIEDITLHEILEYVSPYELERFENEQFAEEREALAVALAAAEAEEERRRQRRKE
C.zeina           82 PG--KLPVFKLKVGDVELEDIGLHEILEYVSPSELERFEHEQFAEEREALAAALAAAEAEDERRRQRRKE
C.zeae-maydis     82 PG--KLPVFKLKVGDVEIEGIGLHEILDYVSPLELERFEHEQFEEEREALAAALAAAAADDERRRQRRKE
```

```
H.werneckii     125 R-ALRKGTIEYQEID----DAGNSGNESYATAGQGRARPTYKNFFKPI------KERRRRKRDPQTGELM
T.nubilosa      135 R-ANTKGVLMRVDPDDH--DAVDSEAFNVATGKHGRARPSYKAQFKML------KARRRRKRNPVTGELM
B.panamericana  132 -RAKLKGVVGYSDDRSELDVGASGEGSEAPAGRHGRARPSYKQLFKKF------KERRRRRRNPVTGELM
F.endolithicus  129 RKAKSNVDARHDDGTGE--PGGDGSAAEVSKGRHGRARPTYKHLYPEL------EKRRRRKRDPRTNELM
F.simplex       146 R-ARLKGVAGYHDGTGE--PGGDEYVAGVIEGKHGRARPNYKHLFKKL------KERRKRKRNPITRELM
R.collo-cygni    60 K-IRKKGIVYFEERNED---LESNEEGEVVLGRHGRARPTYSHLYKVPGGGEDGGKRRRRRRDPSTGDLM
Z.tritici       145 R-ARRKGVVLQVDSGAD---SGADDEGEVAAGRHGRARPTYKHLFKVPEQ----GKRRRRRRDPATGELM
Z.brevis        148 R-ARRKGVVLQVDSGAD---SGADDEGEVAAGRHGRARPTYKHLFKVPEQ----GKRRRRRRDPATGELM
Zas.cellare     131 R-AKTKGQVIYQTDSGT----VSQEDEDVETGKHGRARPTYKHLFKAPEMR---QRKRRRKRDPFTGELM
D.septosporum   132 R-AQHKGIAIFQDLYED--SMEV-DAESVDAGKYGRARPTYKHLFKKM------KERRRRRRDPLTGELM
P.eumusae       261 K-AKQKGTFLFPGVGGD--EDESSDEDDVPQGRRGRARPTYTHLFNRPELP--DERRRRRKRDPETGELM
S.musiva        150 R-TKQKVVPQFEEID-I--DIDEADH-NGSAGGAGRARPTYKHLFQQL-----QKPRRRRKRDPITNELL
C.berteroae     148 R-ARRKGIVLFEEVDDE--DLELADA-NDVVGRHGRARPTYKHLFKVP-----QERRRRRKRDPETGELL
C.beticola      148 K-ARRKGIVLFEEVDDE--ELDSADP-NEVVGRHGRARPTYKHLFKKP-----QERRRRRKRDPETGELL
C.zeina         150 K-ARSKGVVLSEEVNDE--NVDSADP-NEVVGRHGRARPTYKHLFKLP-----EQRRRRRKRDPETGELL
C.zeae-maydis   150 K-ARRKGIVIFEEVNDE--DVDSADP-NEVVGRHGRARPTYKHLFKLP-----EQRRRRRKRDPETGELL

H.werneckii     184 PLSDEESADASD------------------------------------------------------DSLT
T.nubilosa      196 PLSDEDENDDPA------------------------------------------------------ALES
B.panamericana  195 PLSDAEDDALEL-------------------------------------------------------ESS
F.endolithicus  191 PLSDEEIDEAAN------------------------------------------------------TQSS
F.simplex       207 PLSNEEQDVEAN-------------------------------------------------------QSS
R.collo-cygni   126 PLSDEENLHTMGSSDEGVDLNRSSGRSGVRQAPALTSGLQPPKRRRRKRHPITNELMPLSGDENAGSSDG
Z.tritici       207 PLSDEEGGMDV-------------------------------------------------------AASS
Z.brevis        210 PLSDEEGGVDI-------------------------------------------------------AASS
Zas.cellare     193 PLSDEEDAAFN--------------------------A----------------------------PA--
D.septosporum   192 LLSGDENEEAA--------------------------D---------------------------LESSS
P.eumusae       326 PLSDEDLPRPS--------------------------DRGPKRRRRRKRDPLTGELMPLSDHGEDIEMRA
S.musiva        210 PLSDEEVPDS-----------------------------------------------------GDTGMKS
C.berteroae     209 PLSDEDDQMDT---------------------------------------------------S--IAMDS
C.beticola      209 PLSDEDDQMDT---------------------------------------------------C--VPMGS
C.zeina         211 PLSDEDDQVDA---------------------------------------------------GATAAMES
C.zeae-maydis   211 PLSDADDQVDT---------------------------------------------------GATFAMEM

H.werneckii     200 SV----------------------------------------------Q--ARGLQKPSSELLQLQQPTR
T.nubilosa      212 SD----------------------------------------------DNGRPSGSVRLNATGMPEPPKR
B.panamericana  210 AD----------------------------------------------DEPQATL-SSLASSAPAEQHKR
F.endolithicus  207 GD----------------------------------------------DSPARTAPLDERRPAPGDEEKR
F.simplex       222 RD----------------------------------------------DSPAPAAPSREPIRAQTDEQKR
R.collo-cygni   196 GPILQSRTSSSVPIQAAGPSYAQTAEPSMRSNSIFEDSGKPTSAPMQQATIVPSPARRMLPNVTVELPKR
Z.tritici       222 SE---------------------------------E------------EPSRDRSRSAADLLPGVEPHRR
Z.brevis        225 SE---------------------------------E------------EPSRDRSRPAADLLPGVEPHRR
Zas.cellare     207 ---------------------------------------------------KATAASSRAP-APATELPKR
D.septosporum   209 GG---------------------------------E------------QAAMPSS----THMGLPEPAKR
P.eumusae       370 SE---------------------------------E--------------DATTPAARSPAPEAMQLPKR
S.musiva        227 SE---------------------------------E--------------GRLPSVSRAPGPAFSELQKR
C.berteroae     226 SD---------------------------------N------------DQELQSSGNREPAYLGGELPKR
C.beticola      226 SD---------------------------------S------------DQELPPSGSRESAYGGGELPKR
C.zeina         230 PE---------------------------------N------------DQELPSPGHRGPPTTHGELPKR
C.zeae-maydis   230 SD---------------------------------N------------DQELPSSGHREPATTTGELPKR

H.werneckii     222 RRRKRDPVTGELMPLPPLDTSENRMA------------------S----GSLQHKKRPRRRRHPITQELM
T.nubilosa      236 RRRKRDPVTGELMPLEPLGSAKGSHG------------------TSKKLLPFEKKKRRRRKRHPLTGELM
B.panamericana  233 RRRKRDPLTGELLPLDILPQRV-----------------------------SSKKKRQRRRRHPLTGELM
F.endolithicus  231 RRRKRDPVTGELIPLAPLPDATM----------------------------GEEKKRRRRKRHPITLQLM
F.simplex       246 RRRKRDLITGELIPLAPLPEATI----------------------------GEEKKRRRRKRHPVTGELM
R.collo-cygni   266 RRRRRDPITGELLPLDPDPQDVAVSENLS---------------SSRFSNTEERQKRPRRRRHPITGELM
Z.tritici       247 RRRKRDPVTGELLPLSPEVKSSSVAKESPLARSEPAIRSAVGSASSNGDFLQFPPKRPRRRRHPQTGELM
Z.brevis        250 RRRKRDPVTGDLLPLSPEVKSFPAAKESPVARSEPAIQSAVRSAASNGDFLQFSAKRPRRRRHPQTGELM
Zas.cellare     226 RRRKRNKITGELLPLSPDGETAQ--------------------------PSIEKGKRPRRRRHPITGELM
D.septosporum   230 RRRKRDPRTGELLPLDPKPQPSSSRSR-----------NQ----PAEELALGVPYKRPRRRRHPVTGELM
P.eumusae       393 RRRRRDPKTGELLPLDATKSM----------------------------EKDSRYKRPRRRRHPITHELM
S.musiva        250 RRRKRDPITGELLPLSPLQESVPQEVPEPRISQSPALARQ----AKEFFTGLRSEKRGRRRRHPVTQELM
C.berteroae     251 RRRKRDPVTGELLPLPPLPHELGQS-PEASVASVP---------KAELLASLDPNKRPRRRRHPITRELM
C.beticola      251 RRRKRDPVTGELLPLPPLPHELGQS-SEASVASVP---------KVELPVSLEPNKRPRRRRHPITRELM
C.zeina         255 RRRKRDPVTGELLALPPLPHEASQS-PAPSVASVT---------NVGSQASIDHRKLPRRKRHPKTGELM
C.zeae-maydis   255 RRRKRDPATGELLPLPPLPHEASQS-PAPSIASMP---------KVESHTSIDHRKLPRRKRHPKTGELM
```

```
H.werneckii     270 PLGWRYDPNEERRPQA-----SGSKDVSPAMHRMSLSQGEPTKRLKLQHSSYSSDESASEDALAA-----
T.nubilosa      288 PLGWRNDPNNPDDRST---TKAAFGAMSPAMQRMSLSQEQGVKRLKLAHQESEDDGGEQG-SV-------
B.panamericana  274 PIGWRYDPDMPPARQESNSVLAGTIPVSPAFDRLSISDEHPAKRARFDSELSSDLPQSDVESQ-------
F.endolithicus  273 PYGWRYEPNGPSVRPTSGTRRLETGSVSPAMNRLSITMEQPAKRLKLGSRSSSGTSPPAQASIPASVGAA
F.simplex       288 PYGWPYEPNGPSVRPPSNTRRTVTGNMSPAMNRLSITSEQPAKRLKMEGQSSSGGSPSAHVIIASSARVA
R.collo-cygni   321 PLGWVYNPNAGAESFE-----STA--NGSSMQSLSIT--RDSKRVKLASEPSSSSLEDPRPAVAPTSAAS
Z.tritici       317 PLGWRYEDEAERKAWAPKPA-GIESGPSSSMERLSIGNERGSKRIKLTSEPSSGDDRRARPLPFTSPTQE
Z.brevis        320 PLGWRYEDEAEGKACAPKPA-GIESGPSSSMERLSIGNERGSKRIKLTSEPSSGDDRQARPFPSTSPTQE
Zas.cellare     270 PLGWRYDPNTEQQSYEQRRD-GAGAHSTPSIKRLSITREHDAKRMKLTSEEPSSDDELGTRQPVVQ----
D.septosporum   285 PLGWRYEPDVEGNRSE---S-GAGNTSVESMRRLSIVREPELKRPRLASASVASSDPRAQSIDPL---RS
P.eumusae       435 PLGWRYDPDEEKKASP-----SARDLVSPAMRKMSISREHGSKRIKLASESSDDRSQGR--PLPSSSVRM
S.musiva        316 PLGWKYDPSAENESHD-----GRHDGMSPSIERLSISHEQQPKRVKLASEYSADDISREYRSMSPSSRK-
C.berteroae     311 PLGWRYDPETDHV--A-----RKAAGLSPSIRRLSISQEQQPKRVKLASESSADDHPRSANTIRNSPRHA
C.beticola      311 PLGWRYNPETDPV--A-----RKAAGLSPSIRRLSISQEQQPKRVKLASESSADDHTRSANTTRNSPHQA
C.zeina         315 PLGLRYDPKSDTV--V-----RKAAGMKSSMRRLSISQEQRPKRVKLASDSSADTSPLVQATLK-S----
C.zeae-maydis   315 PLGFRYDPKTDTV--V-----RKAAGINPSMRRLSISQEQLPKRVKLASESSADSSPPVHATLK-S----

H.werneckii     330 --------------------------RDP----S-------TEAGTVVTKSPTAQAH-------------
T.nubilosa      347 ----------------------SESDADD----T-------DEVASAATPASALKAV-------------
B.panamericana  337 ----------------------EASEAMP----A-------EQQLTISDVKTPPKPL-------------
F.endolithicus  343 H----------------SDTTSSSEDQP----D-------LK-SWMARSQSSAKQI-------------
F.simplex       358 S----------------SDSTSSSEEEP----A-------LK-SWAAPPQSTPRLP-------------
R.collo-cygni   382 P-----------NVRQPNVIELSESDQEE--QD--DSS--SMQLSIDSAPASKPTTQVRSPKHGM-----
Z.tritici       386 ETTLTPKM-----QATVQVMDLGSEDEDDLAQD--GRTSAQLTPKLSSRPKPSSESK------------I
Z.brevis        389 ETITTPKM-----QTAVQVMDLGLEDEDDLAQD--GRTSAQLTPKLSSRPKPSSESK------------I
Zas.cellare     335 ---------------SRGKLGIFA--SDDS---RESSSSRDPPPMSSSKSRPAPKTSIMNPVSQRISRLG
D.septosporum   348 FLNDGEKQSTAPPKSSDIINELNS--SDS------DDSEDQLQRFSPAQPIPSSKTSNMQPNHKSALSTA
P.eumusae       498 PLRGTPAKSRPIPSAAKVPIDTSE--SDDLARDSENH-----HPISASKPQATPKTSILHPTKAETS---
S.musiva        380 SLSGIPKL-----DNQLVASTIEA--LD---D-------DRAQALRPGMLKSPPKTSMLNPTAQS-----
C.berteroae     374 SPRGTPRQ-----QTKPIVIDLSE--TDDLNDEREKQGKVIPAQ-SPGVTRSTPRTSMLHPTAHT-----
C.beticola      374 SPRGTPRQ-----QQKTIVIDLSE--TDDLNDEKEEQSKAISAP-SPGVTRPTPRTSMLHPTAHT-----
C.zeina         373 PKHGSPHQ-----ETKSIVIDLSE--TDESHDNNEQQHGTRQTQSPPSIPKSTSKTSMLRPTAPT-----
C.zeae-maydis   373 PKQALPHK-----QTKSIVIDLSE--TDDSHANHEQQHGTWQTQS-PSMPKPTPKTSMLRPVVKT-----

H.werneckii     350 --AKSPKSQLGTAAG------AFQ----------------EDASKLRR----YKQTTLTSIMQPTANLSS
T.nubilosa      371 --PSPSQGRLRTPLALQQAAASSTGDTS----P----EPVERKKS-----------TMTSIMQPVAAVTS
B.panamericana  361 --LSASRGQLGVPAALQSAATSSAPETS----P----EPYHGTSLQRR----LIAANTD-------IDNS
F.endolithicus  371 --HSASKGRLGVPVVLQSMATSSAPDTS----P----EPDFPRSAMQT----LGAGNPSPIVQLAVPDSE
F.simplex       386 --PSASKGKLSLPAMLQSTVTSFAPESS----S----EPDLAKSLIRS----PGADNTTSMMQPAALESP
R.collo-cygni   430 MQRARSYSSSNEPITLQSFLSTTATGKAGRDDDETSEDTTSGEETA----LKKTPSQKTSIMNPTASQEI
Z.tritici       437 PLRRSNTSSSEEQITLASFLSGPSKKQESTSDSD---SSLSRRPGKNGANSQKPTPLKNSMMNSQASQGN
Z.brevis        440 PLRRSSASSSEEQITLASFLSGPSKKQESATDSD---SSLSRRPAKSAANSQKPTPMKNSMMNSQASQG-
Zas.cellare     385 PPPSATDSSSEPKITLASFLKPNQPEEDDESSDD---SSSLPKQKVNT----PGERGKTSIMNPMAAQTV
D.septosporum   410 AILSRVDSS-PEPISLASFLKSSKTPGNQEASSDDETDSSSSGPRVKG----TPANGKTSIMNPTGSQTV
P.eumusae       558 -ADDSSDSS-AEPITLASFLKASGRRTHSDDESE--DSEQVAKP--------NRGQVKTSIMNPMAPSQA
S.musiva        428 ---SSSESS-TEPVTLASFLKSSAIQQDQSDSDD--SWSDRQKPIMRK----TMIGGKTSIMNPVATEKR
C.berteroae     431 ---ASSGSS-GERVTVASFLKASATLSDESDTDD--SELFKQKPAVRN----TPTHGRTSILNPIASKQA
C.beticola      431 ---ASSGSS-SERVTVASFLKASATLSDESDTDD--SELFKQKPAVRN----TPTHGKTSILNPIASKQA
C.zeina         431 ---ASSGSA-DEPFTLASFLKASATLADESDTDE--SEVFVKKPGVRS----TPTHGKTSILNPIAAKQS
C.zeae-maydis   430 ---APPGSG-VEPVTLASFLKASAALTDESDSDD--SEVFSKKPTMRR----TPTHGKTSILNPIAAKQP

                                                      CD1
H.werneckii     392 VDDS---GNEDEADDVEDEFPADEYVMEAVLAHHMSDPKTHP---GKEPVMLYQVKWEGYDEPTWEPAES
T.nubilosa      420 AEQS-------------EQDDDEEWIVEDILSHRMSDPRTHPPDLGTKLVLLYHVKWEGDPKPTWEPAES
B.panamericana  410 EGHS-------------AESEEGEFNIDAILDHKLSDPRTHAPEHGKKAVMLYLVKWEGYDEPSWEPVES
F.endolithicus  427 G------------------EESDEWAIEAILGHHMSDPRTHPREFGKKAVMLYQVKWEGFDETSWEPVES
F.simplex       442 DEES-------------DEPGDGEWAIEAILAHHMSDPRTHPRELGNKAVMLYQVKWEGFDEPSWEPVES
R.collo-cygni   496 ADDEDDD------QDIDDELDEGEYFVEAIQAHHMSDPRTHP---GKERVMLYHTKWEGWDEMTWEPAAS
Z.tritici       504 AGGDESE-----SEVSGDNSDDGEWTVEAIVAHHMSDPKTHP---GKPATMLYLTKWEGFPEETWEPAES
Z.brevis        506 AGDDESE-----SEVSGDNSDDGEWTVEAIVEHHMSDPKTHP---GKPATMLYLTKWEGFPEETWEPAES
Zas.cellare     448 EESD---------DAEDEELEEGEWFVEAILGHHLSDPRTHP---GKKQVMLYKTKWEGWNNPTWEPEDS
D.septosporum   475 KPAE--------EDSDSDDLDEDEYVVESILDHHMSDPRSHP---GKSPIMLYKTKWEGFTEPTWEPAES
P.eumusae       616 AE----------VESDEEELEDDEWIVEAIVGHRMSDPNSHP---GRKQVVLYHTKWEGSEEMTWEPAES
S.musiva        488 VTDDDDDDDDDDNGNDDSDLEEGEWFVEAIVGHTLSDPRTHP---GKPSVMLYHTKWEGHQDLTWEPAES
C.berteroae     491 VRPAEDEA----ADKYESHLDDGEWFVEAIVGHKMSDPKTHP---GRPSIMLYKTKWEGHDEPTWEPADS
C.beticola      491 VHPAEDEA----ADNDESDLDDGEWFVEAIVGHKMSDPRSHP---GRPSIMLYKTKWEGHDEPTWEPADS
C.zeina         491 VRSE-EEA----ADDDESDPDEGEWFVEAIVGHRMSDPRSHP---GRPSVVLYQTKWEGHDEPTWEPADS
C.zeae-maydis   490 VHSK-EEA----VDLSGSDLEDGEWFVEAIVRHKMSDPRSHP---DRPSVVLYHTKWEGHDELTWEPADS
```

```
                     CD1
H.werneckii      456 FPDPKTVEEYHRAVKARKPDQPGSSSAGNGLAS--------------KPLPQPGSKRATAP----D-SP-
T.nubilosa       477 FSDPSIIEAYHRMRKQSSRANTASSSQQA----------------RVGSTTAPVNSLMQGT---------
B.panamericana   467 FPDRSVVEHYHGMLKGRQISDERAEEANIDGASPKLVRSSKADSRTSLSLSAPAPK---------H-SS-
F.endolithicus   479 FPDRSVVEEYEREIGARAAIATGTSQNHG--------RGTQAVAAPSVALPNPVNRLLQPAVEPVA-AT-
F.simplex        499 FPDRSVVEEYERRVTAPAAVG-------------------------------------------------
R.collo-cygni    557 FPDQSVVEEYRRRAGLKP----------------------------------------------------
Z.tritici        566 FD-ESIIKDYRRRVGLDKPIGSSNVPKAK---SPANNSPL-------QA--VSAR----AAKSKSPAKFA
Z.brevis         568 FD-ESIIKDYRRRVGLDKPIQSSTASTSK---SPAKNSPL-------QA--VPAR----AAKSASPAKSA
Zas.cellare      506 FVDQGVLEDYRQRAGLNGAKTQIVAG------------SK-------KKSVTP---------APKP----
D.septosporum    534 FPDSSVIDSYGKHVGLKLPEQNRRSTLVT--------GMK--------KTEAPSKAK-VTTRLTSATARV
P.eumusae        673 FLDSSTVRDYHKRLEEFKRNAQSANG--------KVERSK-------QPLGV---AQRETTRVPVP--SI
S.musiva         555 FSDPSTITDYRKRVGLDRDKQASRSPTIV---VPVVASGS-------TSQSALATKGSTASTTKPP--TV
C.berteroae      554 FVDQNTVSEYRARAGLDNPKQVIKSPGRI---VPVVASSS-------RSLTGTSLAR--GPKLRG---S-
C.beticola       554 FVDQNTVSEYRARAGLDNPKQVIKSPGRI---VPVVASSS-------RSLAGPSLAQ--GSKLPG---S-
C.zeina          553 FVDPSTVAEYRARVGLDKAKQTIRSPERI---VPVVASSS-------KPLALPRSSDAQASQLRK---S-
C.zeae-maydis    552 FVDPNTVAEYRARVGLDKAKQAIKSPDRV---VPIVASSS-------KPLTLPKNLDARASQHTK---S-

                                                          CD2
H.werneckii      506 ----VSQHTDGSDDGEEESDEDGIFEIERIVGHRMSDPEEQGLDFGRAPVQLYQVKWRGFEA--LTWETI
T.nubilosa       522 ----VGSR-THPLDLTEEDDEEEGYEIEAILAHELSDPRTHAPDLGREAVMLYKVKWKGYQN--PTWEPV
B.panamericana   526 ----PPQA-MHDAQDSEEEVDDGSFEIEAIVAHHMSDPRTHRPEYGKHPVMLYQVKWKGYNN--TTWEPI
F.endolithicus   539 ----VESS-DDDSDPEDAGSDDGTYIVEAITAHHLSDPQTHAPEFGKKPVMLYKVKWEGYEG--YTWEPT
F.simplex        520 ---------------------------------------------------TVKWKGYEG--HTWEPI
R.collo-cygni        ----------------------------------------------------------------------
Z.tritici        619 QQSEQKPSPTKNMVEIISSDEESWYEVEAILGHHMSDPRTHP---GKPATMLYKVKWVGFKQ--TTWEPS
Z.brevis         621 QQSGQKPSPTKNMVEIISSDEDSGYEVEAILGHHMSDPRTHP---GKPATMLYKVKWVGFKQ--TTWEPA
Zas.cellare      544 ------KATAVNDESDDESDEEEGYEVESILGHHMSDPRTHP---GKPATMLYKVKWLGWPE--PTWEPR
D.septosporum    587 NQ-KGTSVSEEDDDEYNDDDDKVTYEIERILAHHKSDPRTHP---GKPIAMLYQTKWKGYPD--PTWEPA
P.eumusae        723 A---QSKKPVDSSEEETGEDDEEGWEIESILAHHMSDPRTHP---GKPQTMLYKVKWAGWKE--HTWEPL
S.musiva         613 GF-SGNKQQKMNVEEEISESEDEGWEIEKILAHHMSDPRTHP---GKPQTMLYRVKWAGFDESQTTWEPK
C.berteroae      608 -----NSASGSSGNGSEDDGDEDGYEIETILAHHMSDPRTHP---GKPQTMLYKVKWVGYPATAATWEPK
C.beticola       608 -----NSASTSSDGGSEEDEDEDGYEIEEILAHHMSDPRTHP---GKPQTMLYKVKWVGYPASAATWEPK
C.zeina          609 -----KSSSE--SSEDDGEEGQDGYEIEKILAHHMSDPRTHP---GKPQTMLYKVKWVGYPATAATWEPK
C.zeae-maydis    608 -----KSSSG--SLEVDSEGDHDGYEIEKILAHHMSDPKTHP---GKPQTMLYKVKWVGYPATAATWEPK

                          CD2
H.werneckii      570 DSFEDLDVIHDYRRQAGLPALSNDGRIDGCDVCA------------------------------------
T.nubilosa       585 GSFDDLDVVNEYRVRVGLPELEGE----------------------------------------------
B.panamericana   589 ESFEDLSIVNDYRSQAGLPIRDA-----------------------------------------------
F.endolithicus   602 DVFEDRSVVREYRARVGLHAEED-------VAMT------------------------------------
F.simplex        535 DSFEERSVGLPPNKPCPAPDYSEAFPPSRRYVLAELPADALKAPGESAKKLSAK---------VPDYGKR
R.collo-cygni        ----------------------------------------------------------------------
Z.tritici        684 PSLSSTTLLKQYQKRVGLTK--------------------------------------------------
Z.brevis         686 PSLSSTTLLKQYQKRVGLTK--------------------------------------------------
Zas.cellare      603 RSFTDKTIVRKYEKRVGLKPERSEDSNGDMRMKT------------------------------------
D.septosporum    651 NSFTDRKMLHEYQRRVGLRA--------------------------------------------------
P.eumusae        785 ASFPDKSVVNAYRRKVGLIPVKLGANSNLQSSEATVQMEQLNPTTSTHDNTTAHYFGASEAAQTTVGDGN
S.musiva         679 ESFPDVDILNAYRRRVGLKPEPRQ----------------------------------------------
C.berteroae      670 GSFPNLDIVNAYRRKVGLKPETR-----------------------------------------------
C.beticola       670 GSFPNLDIVNAYRRKVGLKPERADCGQ-------------------------------------------
C.zeina          669 GSFPNLDFVNAYRSKVGLEPEKR-----------------------------------------------
C.zeae-maydis    668 GSFPNLDVVNAYRSKVGLKPETR-----------------------------------------------
```

**Figure S5 Sequence alignment of Cbx2 homologs.** Putative Cbx2 homologs from the indicated organisms were aligned using CLUSTAL. Full shading (black) represents conservation of an amino acid in at least 50% of the sequences whilst grey shading denotes conservation of a residue of similar chemistry in at least 50% of the analysed sequences. The location of the chromodomains (CD1 and CD2) is indicated.

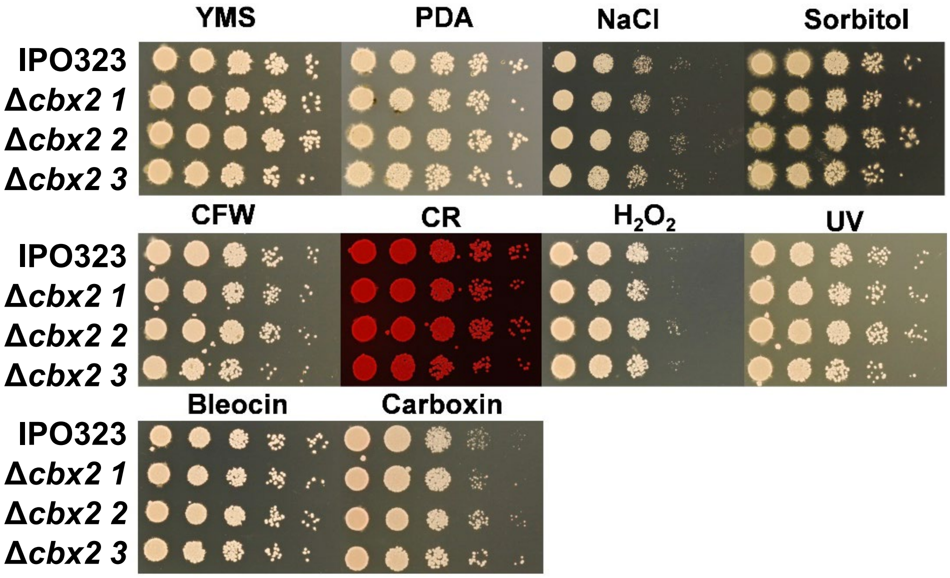


**Figure S6. Deletion of *cbx2* does not result in any detectable reduction in fitness or stress tolerance *in vitro*.** Cell suspensions of the indicated strain were subject to a five-fold dilution series and pinned onto the indicated agar plate and incubated for ~7 days at 18°C. No obvious fitness defect or reduction in tolerance to stress-inducing compounds was observed for the Δ*cbx2* strains. Agar plates were made with YMS (Yeast extract, malt extract, sucrose) or where indicated, PDA (potato dextrose agar). Concentrations of the stress-inducing agents were, NaCl 1 M, sorbitol 1 M, calcofluor white (CFW) 50 µg/mL, congo red (CR) 150 µg/mL, $H_2O_2$ 2 mM, UV dose 250 J/$m^2$, Bleocin 250 ng/mL and Carboxin 2.5 ng/mL.
