## Supplementary Table 5 for "The chromodomain proteins, Cbx1 and Cbx2 have distinct roles in the regulation of heterochromatin and virulence in the fungal wheat pathogen, *Zymoseptoria tritici*"

**Table S5. Oligonucleotide primers used for qPCR in this study**

| **Primer Name** | **Sequence** | **Genomic location** |
| --- | --- | --- |
| qAct1F | TCGTGATTTGACCGACTAC | [10:785364-785382](https://fungi.ensembl.org/Zymoseptoria_tritici/Location/View?r=10:785363-785382;tl=Ce4wnk2eb0Pran0Q-20817104-2059099732) |
| qAct1R | GGATCTCCTGCTCAAAGTC | [10:785246-785264](https://fungi.ensembl.org/Zymoseptoria_tritici/Location/View?r=10:785245-785264;tl=OBPAfmFms9f1B3XX-20817113-2059099985) |
| qGAPDHF3 | GACTACATCGTCGAGTCCAC | [2:1141695-1141714](https://fungi.ensembl.org/Zymoseptoria_tritici/Location/View?r=2:1141694-1141714;tl=O5yDWIIqlqDIrN5i-20817121-2059100289) |
| qGAPDHR3 | GGAGATGACGACCTTCTTC | [2:1141625-1141643](https://fungi.ensembl.org/Zymoseptoria_tritici/Location/View?r=2:1141624-1141643;tl=5HaQ23sVRiQechap-20817128-2059100342) |
| qtelomere00029F | CGCTCTCGGTAACCTAGC | 1:169707-169724 |
| qtelomere00029B | GCGCTAGGCTATACGGAC | 1:169783-169800 |
| qDTHelementF | GCCGCCGCAGCTGTATAG | 9:29043-29060 |
| qDTHelementB | CTACCCTACGCCTGCGAC | 9:29109-29126 |
| qMycgr3G103556F | GACGTTCTCCTCTTGTGCAG | 3:409287-410075 |
| qMycgr3G103556B | TGTCATGCTTTCGGGAACAC | 3:409287-410075 |
| qMycgr3G44980F | ACTACTCCGATGCTCTGACC | 7:2607080-2608068 |
| qMycgr3G44980B | TTTCAGATTCGGGCACACTG | 7:2607080-2608068 |
